# Immune phase transition under steroid treatment

**DOI:** 10.1101/2021.01.19.427269

**Authors:** Sonali Priyadarshini Nayak, Susmita Roy

## Abstract

The steroid hormone, Glucocorticoid (GC) is a well-known immunosuppressant that controls T cell-mediated adaptive immune response. In this work, we have developed a minimal kinetic network model of T-cell regulation connecting relevant experimental and clinical studies to quantitatively understand the long-term effects of GC on pro-inflammatory T-cell (T_pro_) and anti-inflammatory T-cell (T_anti_) dynamics. Due to the antagonistic relation between these two types of T-cells, their long-term steady-state population ratio helps us to characterize three classified immune-regulations: (i) weak ([T_pro_]>[T_anti_]); (ii) strong ([T_pro_]<[T_anti_]), and (iii) moderate ([T_pro_] ∼ [T_anti_]); holding the characteristic bistability). In addition to the differences in their long-term steady-state outcome, each immune-regulation shows distinct dynamical phases. In the pre-steady, a characteristic intermediate stationary phase is observed to develop only in the moderate regulation regime. In the medicinal field, the resting time in this stationary phase is distinguished as a clinical latent period. GC dose-dependent steady-state analysis shows an optimal level of GC to drive a phase-transition from the weak/auto-immune prone to the moderate regulation regime. Subsequently, the pre-steady state clinical latent period tends to diverge near that optimal GC level where [T_pro_]: [T_anti_] is highly balanced. The GC-optimized elongated stationary phase explains the rationale behind the requirement of long-term immune diagnostics, especially when long-term GC-based chemotherapeutics and other immunosuppressive drugs are administrated. Moreover, our study reveals GC sensitivity of clinical latent period which might serve as an early warning signal in the diagnosis of different immune phases and determining immune phase-wise steroid treatment.

## I. Introduction

The dynamics of biological regulatory networks, their adaptation under different environmental stresses, how they are misguided and diseased are all timely and relevant global questions [1–4]. For instance, in the current pandemic situation, our utmost focus lies on the human immune network, which undertakes a cascade of cellular interaction and biomolecular reactions to protect us against a universe of pathogenic microbes. The human immune system is a highly complex network where the immune cells are in continuous interactions and clashes with foreign invaders/pathogens to maintain a healthy state. One of the key targets of this immune system is to distinguish between self-cells and non-self-cells. In the consideration of its way of operation, the immune response has two interconnected arms in the form of two subsystems, i.e., innate immunity and adaptive immunity. While the innate immune system is a non-specific type of defense mechanism which is present in our body from the time of birth, adaptive immunity is a subsystem of the immune system which comprises specialized, systemic cells but a slow pace response process. Among different lymphocyte populations of the adaptive immune system, although CD4+ T-cells play a significant role in the immune responses throughout the defense mechanism against the pathogen, on the contrary, some pro-inflammatory CD4+ T-cells often fails to distinguish between self and non-self-cell, causing some auto-immune diseases and allergies. Among these CD4+ T cells, some act as pro-inflammatory cells, others as anti-inflammatory cells. The regulatory/anti-inflammatory T cells exert a down-regulation mechanism on the population of effector/pro-inflammatory T cells to prevent auto-immunity [5–11].

The human body has a myriad of feedback loops and mechanisms to balance the dynamic equilibrium of the cell populations for the proper functionality of a healthy body. Along with the regulatory anti-inflammatory T cell, secosteroid hormone like Vitamin D and steroid hormone like Glucocorticoid(GC) [12–19] also evolve to supplement its immunomodulatory action. Vitamin D and GC, both downregulate the pro-inflammatory T-cell population and upregulate the anti-inflammatory T cell population [1,2,4,12–14,20–22]. In our early study, we have developed a coarse-grained but general kinetic model in an attempt to capture the immunomodulatory role of vitamin-D to control the population ratio between pro-inflammatory and anti-inflammatory T-cell populations. We revealed a nonlinear effect of vitamin-D on T-cell regulation which is an indirect result of antigen presentation and subsequent production of pro-inflammatory effector T-cells [4]. In subsequent work, borrowing concepts from equilibrium statistical mechanics, we introduced a new description of the immune response function in terms of fluctuations in different subsets of T-cell [3]. We found a divergence-like growth near the co-existence line of distinct immune phases, which is a characteristic of dynamic phase transition. A phase transition phenomena, in general, is coupled to an external perturbation. In our T-cell regulation model, we are focused on deriving the GC dose dependence of T-cell dynamics. Along with that, we also intend to draw a phase diagram delineating different immune phases over a sensitive order-parameter domain.

To envision a multi-dimensional phase diagram distinguishing different immune phases in the field of immunology is a relatively new concept compared to its wide range of applicability in physical chemistry, engineering, mineralogy, and materials science [23–26]. On the other hand, the framework of mathematical models dealing with cellular dynamics that drive the crossover from one phase to the other has long been one of the major topics in cell biology. In such studies, microbial cell growth dynamics are monitored under different environmental conditions. In the case of bacterial cell growth, the environmental drivers are oxygen, pH, temperature, or availability of nutrients, to name a few [27]. In a laboratory, under optimal conditions, a canonical microbial growth curve follows essentially four different phases: (i) lag phase, (ii) log phase, (iii) stationary phase, and (iv) death phase. The exponentially growing log phase has led to the development of several growth laws, while an emerging stationary phase is observed to halt the growth under critical environmental stress. Once cells enter the stationary phase, a certain time-span is generally required to recover growth after the condition tends to renormalize [28–30].

In recent time, the dynamical pattern of CD4+ T cell counts those are HIV infects has been monitored to follow the disease progression. For clinicians, CD4+ T cell count and viral levels in the plasma are the key markers to navigate the disease progression. Also, in such cases, after an acute infection period (2-10 weeks), CD4+ T cells enter a stationary phase clinically termed as ‘clinical latent phase’. This is an apparent near-normal asymptomatic phase, where viral load drops dramatically. However, in this phase, HIV is continuously infecting new cells and actively replicating. After a long asymptomatic period (more than 15 years as evidenced), the virus enters into a resurrection phase and eventually gets out of control to destroy the remaining cells [31,32]. A very recent kinetic model has attempted to characterize the role of GC on the immune system and anti-tumor immune response over 30 days period under a constrained GC supply [33]. However, several early clinical reports suggest that most immunosuppressive and chemotherapeutic GC based drugs at their high dose have rather a long term (in terms of years) effect, and the adverse effect/s of these drugs may arise even long after the treatment has stopped [34–36]. From such above cases, it is evident that long-term immune dynamics under GC administration needs to be studied.

GC drugs are being used in the field of medicine for more than 65 years. Though there are several classes of cost-effective synthetic GCs, Dexamethasone (Dex) is the most widely used because of their higher binding affinity to GC receptors (GR) than natural cortisol; additionally, it has minimal mineralocorticoid activity. However, it is much more potent and has a longer duration of action as compared to other synthetic GC like prednisolone and prednisone [37–40]. GC exert their primary anti-inflammatory and immunosuppressive effects on both innate and adaptive immune response [18,19]. It has been reported in various experimental findings that GC mediates the inhibition in the maturation process of DC via downregulation of CD80/86, CD1a, MHC class II, and reduced cytokine synthesis including IL-12 and TNFα [37,39,41]. In the work of Cook *et al*., they have reported large-scale depletion of lymphocyte, particularly CD4+ T cells and CD8+ T cells but significant increase in the activation and proliferation of regulatory T cell/anti-inflammatory T cell by increasing expression of both Ki67 and ICOS, contributing to their immune suppressive activity. Moreover, they have also recorded change in the Dendritic cell (DC) subtypes population, a similar phenomenon has also been observed in various other *in-vivo* and *in-silico* models [18,33,42–44].

In this current study, to understand the immunomodulatory role of steroids, we have taken into consideration synthetic Glucocorticoids, Dex-mediated immune phase transition of adaptive immune response mainly on the CD4+ cell (pro-inflammatory cells and anti-inflammatory T cells). We have developed an interaction-based kinetic scheme, which is depicted in Fig. 1 to portray the direct and indirect effects of GC on the immune system.

**FIG. 1:**
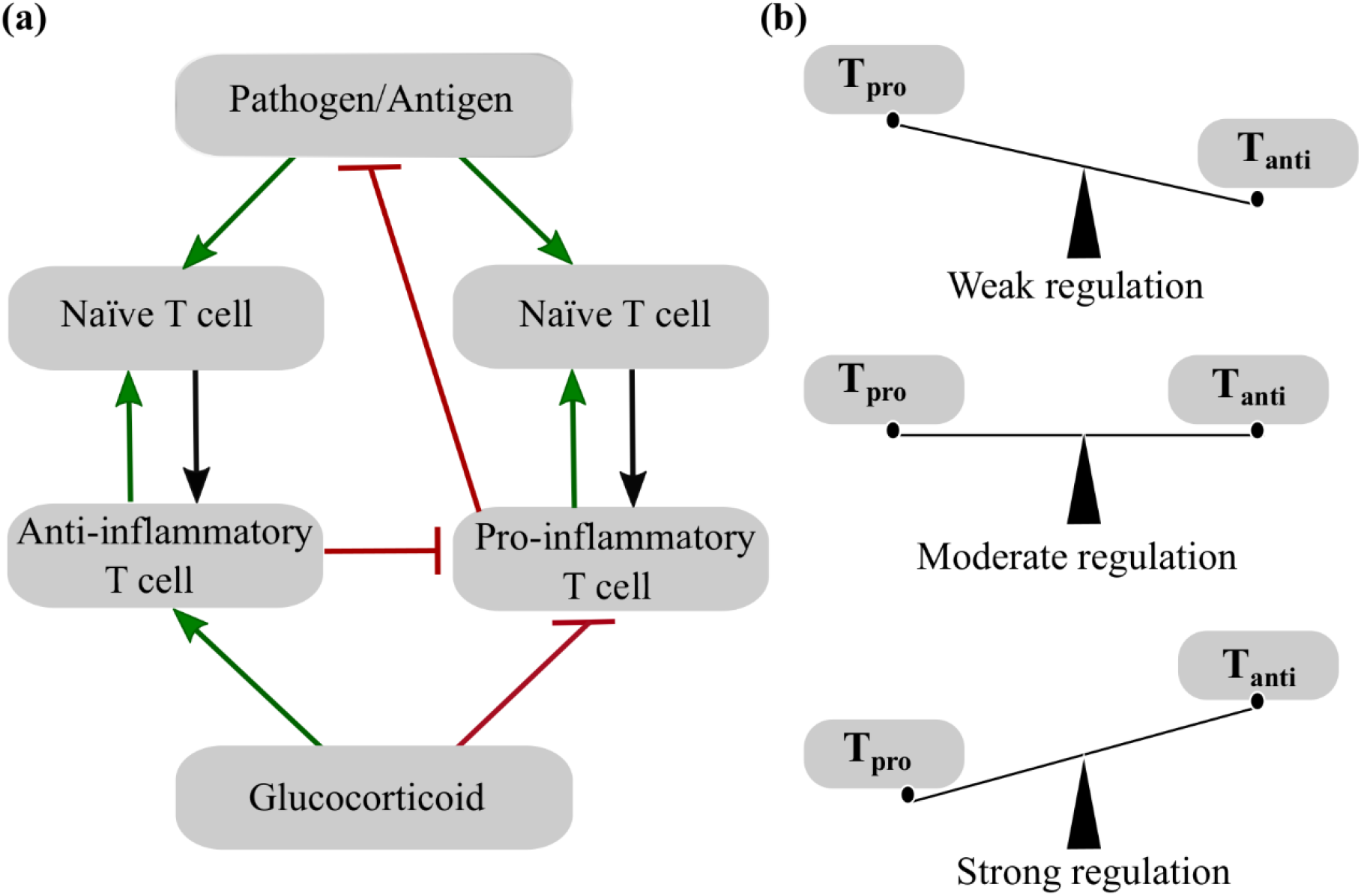
Coarse grain model of the adaptive immune response in the presence of GC. (a)There are five primary elements in our system include pathogen/self-cell containing the antigen, CD4+ Naïve T cell, anti-inflammatory T cell, pro-inflammatory T cell, and Glucocorticoid. The overall interaction among these five elements are presented in the network. Perturbation from the pathogen in the body leads to the activation of Naïve T cells to mature into pro-inflammatory and anti-inflammatory T cells. Mature pro-inflammatory T cells cause the killing of pathogens or self-cells containing the antigen. Further, the mature pro-inflammatory T cells and anti-inflammatory T cells have a role in the self-regulation process of inducing naïve T cells to produce more of themselves, respectively. To control the over-explosion of pro-inflammatory T cells. Anti-inflammatory T cells and GC cause its down-regulation. In the given flowchart, the green arrows stand for the upregulation process, and red arrows for the inhibition process, and the black arrow represents conversion processes. (b) It represents the population balance of pro-inflammatory T cell (T_pro_) and anti-inflammatory T cell (T_anti_) across the three regulations: weak (T_pro_>T_anti_), moderate (T_pro_∼T_anti_), and strong (T_pro_<T_anti_).

## II. Model and Method

The present kinetic immune-network model is developed based on several early experimental and mathematical model studies. Our immune system is comprised of complex and diverse network modules that accompany many participants, which are strongly coupled with each other resulting in synergistic interaction for the maintenance of a healthy physiological condition [1,2,45]. To understand such complex interactions among different immune cells, pathogens, and also to characterize the immunomodulatory role of Glucocorticoid (GC), we need to develop a simple modelling pathway that can be interpreted and explained. To understand the correlation among these different immunological interactions of diverse cell types and pathogens, we have to look carefully at how the cells are coupled and how the immunomodulator affects their overall interaction.

### A. Development of the reaction network model of CD4+ T-cell regulation with and without GC

After careful filteration of all the essential and most important participants, we develop a simpler and refined correlation among the various elements of the immune system, which is presented in Fig. 1. Once the developed network appears to be simple and effective enough, a system of coupled differential equations is used to model the system.

The important participants considered in our coarse-grained model are the following: (i) Pathogen/antigen/self-antigen, (ii) Naïve T cell (precursor T cell), (iii) Pro-inflammatory T cell, (iv) Anti-inflammatory T cell, and (v) Synthetic glucocorticoid (dexamethasone)

To create a simple albeit effective model of these CD4+ T cells regulation and modulation, we perform model analyses based on CD4+ T-cell activation, deactivation, and regulation, following some in-vivo and in-silico results discussed below.

I. We have grouped all the CD4+ T cells which cause inflammation and allergic response by down-regulation of pathogens; those CD4+ T cells are tagged as pro-inflammatory T cells, which include Th 1, Th 2, Th 17, Th 9, Th 22, and Tfh. On the other hand, we grouped all inflammation suppressing or downregulating the role of pro-inflammatory T cells as anti-inflammatory T cells, which include Th3 and Treg. Both anti-inflammatory T cells and pro-inflammatory T cells are the lineages of CD4+ T cells [10,46–53].
II. As these immune cells are in continuous interaction and clash with foreign invaders/pathogens to maintain a healthy state, we can say that there is a continuous predator-prey tussle between pro-inflammatory T cell (effector T cell) and the pathogen, where pro-inflammatory T cell being the predator and on the other end pathogen being the prey [9,54,55].
III. Upon perturbation from pathogens and/or tissue trauma, pattern recognition receptors detect cytokine-induced danger signals. These cytokines induce the production of more of itself through various biological pathways, which result in the amplification of inflammation. GC plays a crucial role in such a situation. Being the immunosuppressant it downstreams the danger sensor by causing inhibition of many pro-inflammatory cytokines expression, which includes granulocyte-macrophage colony-stimulating factor (GM-CSF), interferon-γ(IFNγ), TNF, IL-4, IL-5, IL13, IL-9, IL-17. However, GC also controls the production of cytokine at the post-transcriptional level. It decreases the half-life of TNF mRNA by upregulating tristetraprolin. In this way, GC exerts its anti-inflammatory and immunosuppressive effects on the adaptive immune response. Early studies have reported large-scale depletion of lymphocyte, particularly on CD4+ T cells. However, they have also noted a significant increase in the activation and proliferation of anti-inflammatory T cells by increasing expression, contributing to their immune suppressive activity [18– 20,37,42–44,56,57].
IV. Glucocorticoid (GC) has a modulatory effect on both the population of CD4+ T cells (anti-inflammatory T cells and pro-inflammatory T cells). GC downregulates the pro-inflammatory T cell, i.e., effector T cell population, and on the other hand, they upregulate the anti-inflammatory T cell population. GC maintains a perfect balance between these T cell populations to maintain the homeostasis of the body [20,33].

In the present context, we analyze the following set of biological transformations. Most of them are catalytic reactions in terms of up-regulation or down-regulation.

A. The initial step is the elimination of the pathogen by pro-inflammatory T cells.

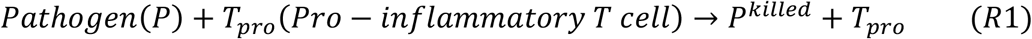
B. Further pathogenic contact and/or pro-inflammatory T cell contact promotes the maturation of naïve(precursor) T cells into mature pro-inflammatory T cells.

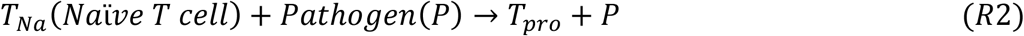

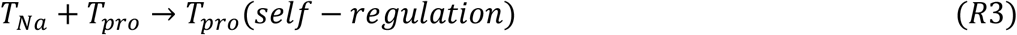
C. Similarly, pathogenic contact and/or anti-inflammatory T cell contact promotes the maturation of naïve(precursor) T cells into mature anti-inflammatory T cells.

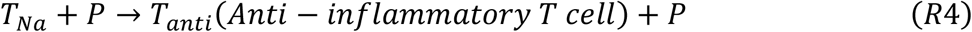

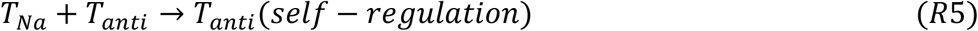
D. Both anti-inflammatory T cell and active Glucocorticoid (GC^*^) can suppress (down-regulate) the pro-inflammatory T cell production to control the hyperactivity of the immune system.

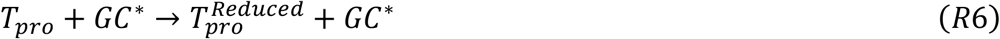

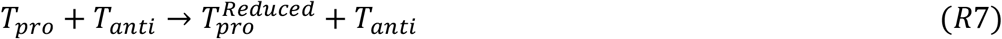
E. On the other hand, GC upregulates the production of anti-inflammatory T cells.

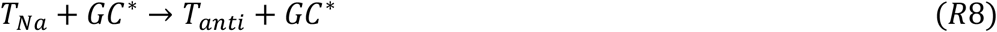

### B. Master equations quantifying the reaction network dynamics

Now, some essential additional presumptions we set before writing the kinetic equations:

- For pathogen and naïve T cells, each has a birth rate, which contains influx and proliferation rates, and death rate similar to the decay, which incorporates scenario of natural cell death. The death rate of all the components is linear with its concentration.
- All the rate parameters are assumed to be constant as obtained from both in-silico and in-vivo models and experiments but it may vary from system to system (i.e., here person to person).
- To scale the unit, here we assume that in the absence of pathogen, a hundred (average number of T-cell present in hundred nano-liter of a blood sample) naïve T-cells [58] pre-exist, which gives a concentration of 0.00000166 nmol/Litre, an elaborate calculation is described in Appendix B.

The above recombination, annihilation, and catalytic reactions lead to the following set of coupled equations. The equations are size-extensive. However, the size extensibility is the critical robustness of our model. We believe that we are not missing any dynamic character of the system. However, the present method we have employed is a deterministic approach, that is, these set of coupled equations are solved in a deterministic way.

### C. Development of two generic models to assess GC induced reaction network dynamics

In this study, we have modelled the effects of GC on different subsets of immune cells considering two different modes of GC’s intake: (i) Model-I: In this model, along with external administration of GC, we have taken into account natural cellular production GC maintaining its pharmacokinetic characteristics. Here, GC induced pro-inflammatory T-cells inhibition is linearly dependent on GC concentration following an early treatment [4]. (ii) Model-II: GC mediated inhibition was done by applying a saturation function of GC concentration as used by Yakimchuk in a very recent work [33] Here, one-time external intake of lower dose of GC and its exponential decay has been considered, for comparison purpose.

### MODEL I

In Model-I, initially, we have considered a system free of GC to get a better understanding of the immune response in the absence of any drug. Couple differential equations for the system in the absence of GC is shown in Appendix A. Five coupled ODEs of Model-I in the presence of GC are presented below.

### Corresponding five kinetic equations for GC regulation

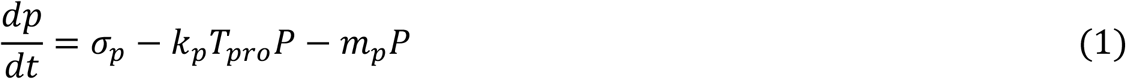

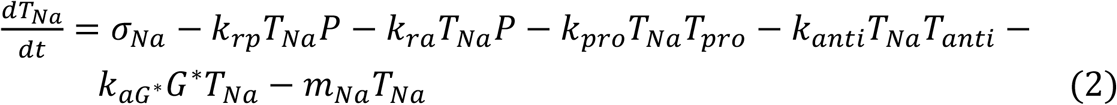

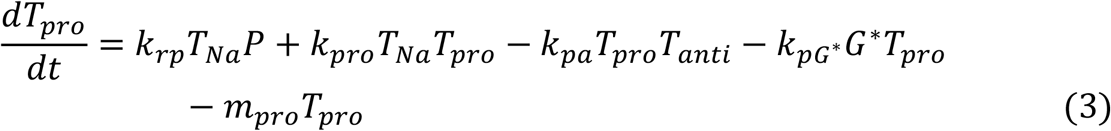

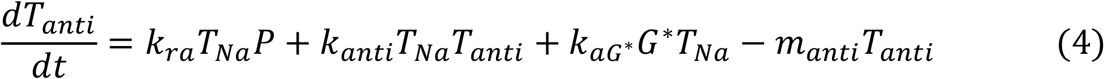

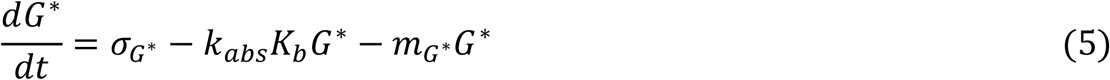

### Model II: Replacing the rate equation of GC by a saturation function

Here, we have followed the method as described by Yakimchuk [33]. to reduce the number of coupled differential equations by considering the concentration change of GC with time using a saturation function. Here also, our kinetic scheme is same as written in the above method part, while only the activated GC concentrations are replaced by a saturation function which takes into account the decay rate of GC. Hence, GC’s pharmacokinetics has not been included here. However, the usage of saturation function has some limitations. It is restricted only to the lower values of GC dose so as to capture all the three regulation regimes. As the [1-exp(-G*)], tends to one with an increase in GC dose, as exp(-G*) tends to zero, left us with a saturated system. Once it is saturated, a further increase in GC dose will have a null effect on the overall system.

### Glucocorticoid Saturation Function(G)

*G* = *k*_*i*_ (1 − *e*^−*G*\*^), *k*_*i*_*=*inhibition rate for a particular immune cell type,

*G* = *k*_*a*_ (1 − *e*^−*G*\*^), *k*_*a*_*=*activation rate for a particular immune cell type

*G*^*\**^= initial glucocorticoid concentration

### Coupled ODE for our system accounting for glucocorticoid concentration as saturation function

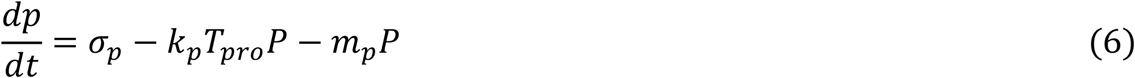

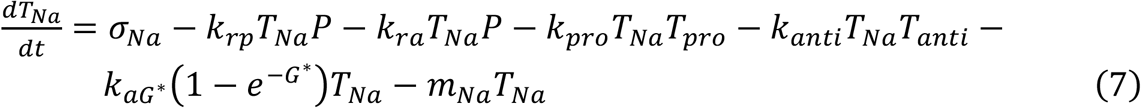

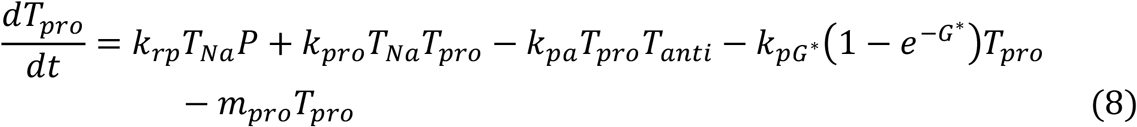

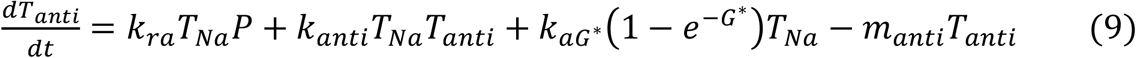

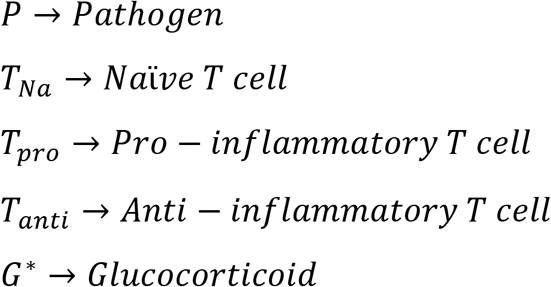

### D. Parameter estimation, steady-state and stability analysis

In order to solve these five coupled differential equations by a deterministic approach, we need to estimate the parameter values associated with these equations. However, the determination of accurate values of all parameters is quite daunting as the rate constants depend on several other factors and differ from one species to another; therefore, there is no universality of the rate constants. So, in order to overcome this problem, we employ diverse approaches for the determination of the rate constants. For some of the cases in which the rate constant has either been reported in the literature or can be calculated from the literature, we have used that value from the literature. In some cases, where the order of magnitude of the rate constants have been reported in the literature, that order is taken as the parameter value. The values of parameters taken for solving the coupled differential equations are listed in Table-I. Here we have used the same formalism as developed by Fouchet *et al*. to obtain and estimate the parameter values [59].

**TABLE 1:** Basic parameter values (*time duration is taken as ‘‘days’’).

| Si No. | Parameter | Symbol | Value |
| --- | --- | --- | --- |
| 1 | Reproduction rate of pathogen | $\sigma_p$ | 1 |
| 2 | Rate of pathogen killing by pro-inflammatory T-cells | $k_p$ | $10^2$ |
| 3 | Death rate of pathogen | $m_p$ | 1 |
| 4 | Birth rate of naïve T-cell | $\sigma_T$ | 1 |
| | Death rate of naïve T cell | $m_{Na}$ | 0.01 |
| 6 | Rate of differentiation of naïve T cell to pro-inflammatory T cell which is induced by antigen | $k_{rp}$ | 1.01 |
| 7 | Rate of differentiation of naïve T cell to pro-inflammatory T cell which is induced by Pro-inflammatory T cell itself (auto catalytic) | $k_{pro}$ | variable |
| 8 | Rate of differentiation of naïve T cell to pro-inflammatory T cell which is induced by Pro-inflammatory T cell itself (auto catalytic) | $k_{anti}$ | $10^{-1}$ |
| 9 | Rate of differentiation of naïve T cell to anti- inflammatory T cell which is induced by pathogen | $k_{ra}$ | 1.01 |
| 10 | Rate of inhibition of pro-inflammatory T cell by anti-inflammatory T cell | $k_{pa}$ | $10^2$ |
| 11 | Rate of decay of pro-inflammatory T cell | $m_{pro}$ | 0.1 |
| 12 | Rate of decay of anti-inflammatory T cell | $m_{anti}$ | 0.1 |
| 13 | Production rate of Glucocorticoid | $\sigma_{G*}$ | 1.872 |
| 14 | Rate of inhibition of pro-inflammatory T cell by active Glucocorticoid(dex) | $k_{pG*}$ | 0.57 |
| 15 | Rate of anti-inflammatory T cell reactivation by active Glucocorticoid(dex) | $k_{aG*}$ | 1.0483 |
| 16 | Absorption rate of Glucocorticoid(dex) | $k_{abs}$ | 102.857 |
| 17 | Bioavailability of Glucocorticoid(dex) | $k_b$ | 0.75 |
| 18 | Rate of decay of Glucocorticoid(dex) | $m_{G*}$ | 0.415 |

The concentration of naïve T-cell is calculated to be 1.66 × 10^−6^ nmol/L in the absence of any antigen/pathogen for detailed calculation (see Appendix B). Moreover, these naïve T-cells have a turnover of 1% per day. The concentration of pathogen has also been normalized by setting the birth and death rate of the pathogen to the same value. This is done so that at the steady-state, the concentration of pathogen will be 1. The decay rate of both pro-inflammatory and anti-inflammatory T-cells are arbitrarily set to the same value, 0.1, as they account for proliferation and death rate altogether for both the subset of T-cells. So, setting them to the same value leads to mutual compensation, and thus equilibrium is not affected much by their values. Apart from birth rates and death rates, other rate constants are taken from various literature [33,59–62], whose detailed estimations are explained in Appendix B, which also includes related pharmacokinetic rates of glucocorticoid (Dex).

We have also done a steady-state and stability analysis of the system. A detailed description is shown in Appendix C.

### E. Classification of T-cell regulation

Based on our early study [3,4], we have classified T-cell regulation into the following three groups: (i) **weak regulation**; where the concentration of pro-inflammatory T cell is very high, and the concentration anti-inflammatory T cell is low, which result in a lowered number of the pathogen, in this phase the immune system is auto-immune prone; (ii) **strong regulation**; where pro-inflammatory T cell concentration is low, which is a result of higher concentration of anti-inflammatory T cell leading to high pathogen population, this can be an immunocompromised condition, where our body is prone to disease, and (iii) **moderate regulation;** where the concentration ratio of pro- and anti-inflammatory T cells are balanced and the immune system holds the characteristics of bistability.

## III. Results and discussion

Immunosuppressive drugs are unavoidably correlated with an increased risk of immuno-compromised conditions with infection and malignancy. Several studies reported that GC concentration is directly proportional to the growth and enhancement of anti-inflammatory T-cell populations and alongside GC suppresses pro-inflammatory T cells. Thus, the dynamics of T-cell are very sensitive to the dose of GC [4,37,42,63,64]. Hence, an optimal level of GC administration is essential for the proper functioning of the human body.

Our immune system is a dynamic network encompassing numerous events with a parameter space which may vary from individual to individual. We have accounted for these events through coupled kinetic rate equations where we have included the values of rate constants and initial pre-existing concentrations of naїve T-cells, and GC concentrations as initial inputs. In some instances, a rate constant or a set of rate constants may show higher sensitivity and variability compared to other rate constants. This can be considered as a person based diversity in the immune system. Thus, we are also interested in exploring sensitive rate parameter/s and how different sets of parameters control the immune response to the invasion by antigens, including dose dependence of GC.

### A. GC treated and untreated CD4+ T cell dynamics and different dynamical phases

We consider a system both with and without GC treatment. By solving the above-mentioned five coupled differential equations of Model I, we find the dynamical behaviour of pro-inflammatory and anti-inflammatory T cells throughout their course of time evolution, as depicted in Fig. 2, in the presence of GC. In the absence of GC, we have only four coupled rate equations described in Appendix A and the corresponding dynamical behaviour of T-cells are shown in Fig. S1 of supplementary material. As several early clinical studies reported that immunosuppressive drugs like GC often leave a long-term effect even long after the treatment has stopped [34–36], in this study, we monitor GC-treated and untreated T-cell dynamics over almost a year-long period. In both cases, after observing the long-term time-evolution of T-cells under small pathogenic perturbation limit, we identify different dynamical phases of T-cells and thus classified majorly into three periods/phases: Expansion period (lag and log phase), latent period (intermediate stationary phase), and finally, a long-term steady-state as shown in Fig. 2. Similar dynamical phases in terms of lag, log, and intermediate stationary states are well-known in the time evolution of microbial growth pattern found in various experimental studies [28,30]. However, in T-cell dynamics studies, this unique *pre-steady state stationary phase* behaviour sustaining for a few months long period in the intermediate time progression range has not been characterized in any early work. Long-time dependent T-cell regulation studies are limited [3,31,32].

**FIG. 2:**
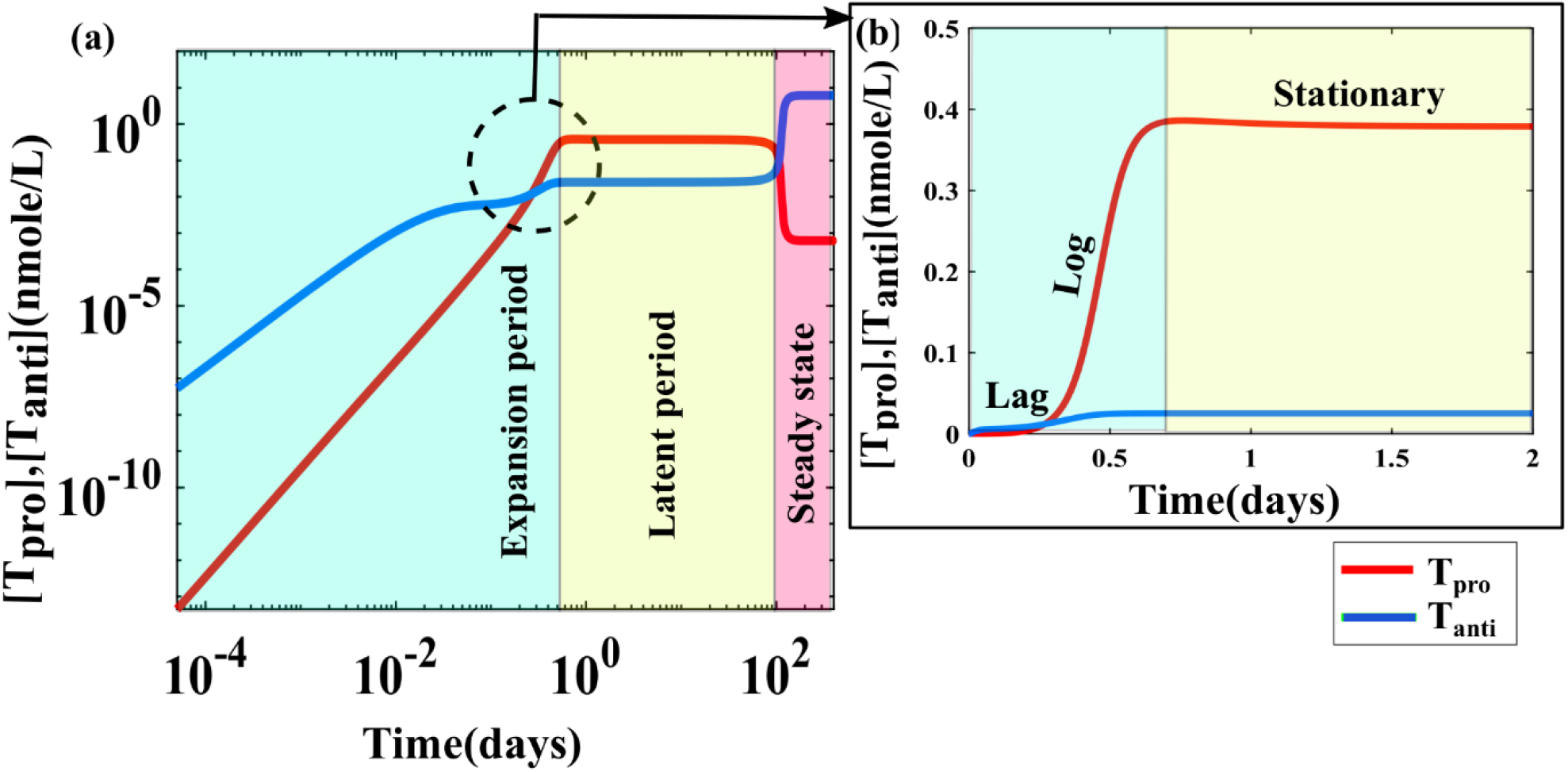
Time evolution of the immune response of CD4+ T cells in the presence of GC. (a)The log-log plot represents a regime of the immune phase regulation that contains an expansion period, followed by a latent period, i.e., the time range within which the concentration of pro-inflammatory T cells and anti-inflammatory T cells does not change with time. After the latent period, there is a jump to the final steady-state condition. (b)Moreover, the schematic illustration of the phases of immune response mediated by antigen-specific pro-inflammatory T cells is depicted on the left side, where three phases of the initial T cells immune response (lag phase, log phase, and stationary/latent period) are indicated. The time evolution of pro-inflammatory T cells and anti-inflammatory T cells are plotted by solving the coupled kinetic equations using a deterministic approach. In the plot, pro-inflammatory T cells and anti-inflammatory T cells are represented by the red line and the blue line, respectively. The cyan shaded region represents the expansion period, the yellow shaded region represents the latent period, and the pink shaded region represents the steady-state. Note that here we consider *k*_*pro*_=56, and the other rate values are the same as given in Table-I. Note the zoomed portion of (a) is shown in (b).

In this stationary phase, the pro-inflammatory and anti-inflammatory T cells maintain highly balanced concentrations, and very less amount of pathogens are observed to present which are not likely to cause any disease-related disorder, as shown in Fig. S2. In clinical terminology, it is considered as an asymptomatic phase and the phase duration as a clinical latent period. A similar latent period is observed in the case of various HIV based models [31,32] containing a period of clinical latency where the patient does not exhibit any symptoms. Finally, at a longer time, the system reaches a steady-state with a higher number of anti-inflammatory T cells and a lower number of pro-inflammatory T-cells in the post latent period depicting the antagonistic nature of pro-inflammatory T cells and anti-inflammatory T cells.

In connection to early time evolution study of CD4+ T-cell [65], here also, we find that the initial phase of pro-inflammatory T cell time evolution has three sub-phases: expansion, contraction, and memory. The pro-inflammatory T cells clonally increase in number during the first phase, in the presence of antigen. Soon after the pathogen load dropped down, the contraction phase follows, and the number of pro-inflammatory T cells reduces due to apoptosis. After the contraction phase, the number of pro-inflammatory T cells stabilizes and is maintained for significant periods, representing the memory phase, as depicted in Fig. S3 of supplementary material. Similar three phases have also been reported in other studies [65].

### B. Effect of Glucocorticoid on CD4+ T-cells population: Transition from weak to moderate to strong regulation

Mature pro-inflammatory T cells are the ones responsible for the elimination of pathogen/malignant self-cells [7,9,66]. However, these pro-inflammatory T cells have a role in the self-regulation process of inducing naïve T cells to produce more of themselves. This phenomenon is evident from various studies where it has been suggested that the pro-inflammatory cytokines produced from mature activated CD4+ T cells induce the production of more of itself through various biological pathways [20,67,68], this over-amplification of inflammation may lead to an auto-immune disorder. In various model studies, it has been reported that the anti-inflammatory T cells maintain a balanced regulation of the immune system [3,4,33,59]. Moreover, numerous clinical and experimental studies suggest that the immunomodulatory role of GC causes down-regulation of the pro-inflammatory T cells population to keep them under control [15]. To observe the role of GC in the interaction network of the immune system, we introduce GC-related rate constants and initial pre-existing GC concentration into our system of consideration. After the introduction of GC, we observed that GC has the potential to modulate the immune system from weak regulation to moderate regulation to strong regulation. Along with GC, we have found another sensitive rate constant *k*_*pro*_, (autocatalytic rate of pro-inflammatory T cells), which also has the ability to modulate the immune system across these three regimes both in the presence and absence of GC.

To investigate several GC associated factors, we have performed time evolution analysis of each participating element after perturbation from pathogen to study their long-time behaviour by varying *k*_*pro*_ both in the absence and presence of GC. By solving our system of equations, we have got all three regulation regimes, both in the presence and absence of GC. Fig. 3(a) shows that in the absence of GC, the system falls under a weak regulation limit when we fix *k*_*pro*_ = 50. However, In the presence of a standard level of GC, we found weak regulation Fig. 3(b) at *k*_*pro*_=70. Fig. 3(c) shows a moderate regulation in the absence of GC at *k*_*pro*_=30, where the intermediate stationary phase sustains with a latent period of 20 days. Furthermore, in the presence of GC, we have found a moderate regulation with stationary phase extended with a latent period of 90 days at *k*_*pro*_=56, Fig. 3(d). In both cases (absence of GC), as shown in Fig. 3(e) and (presence of GC) Fig. 3(f), we find strong regulation at *k*_*pro*_ =10.

**FIG. 3:**
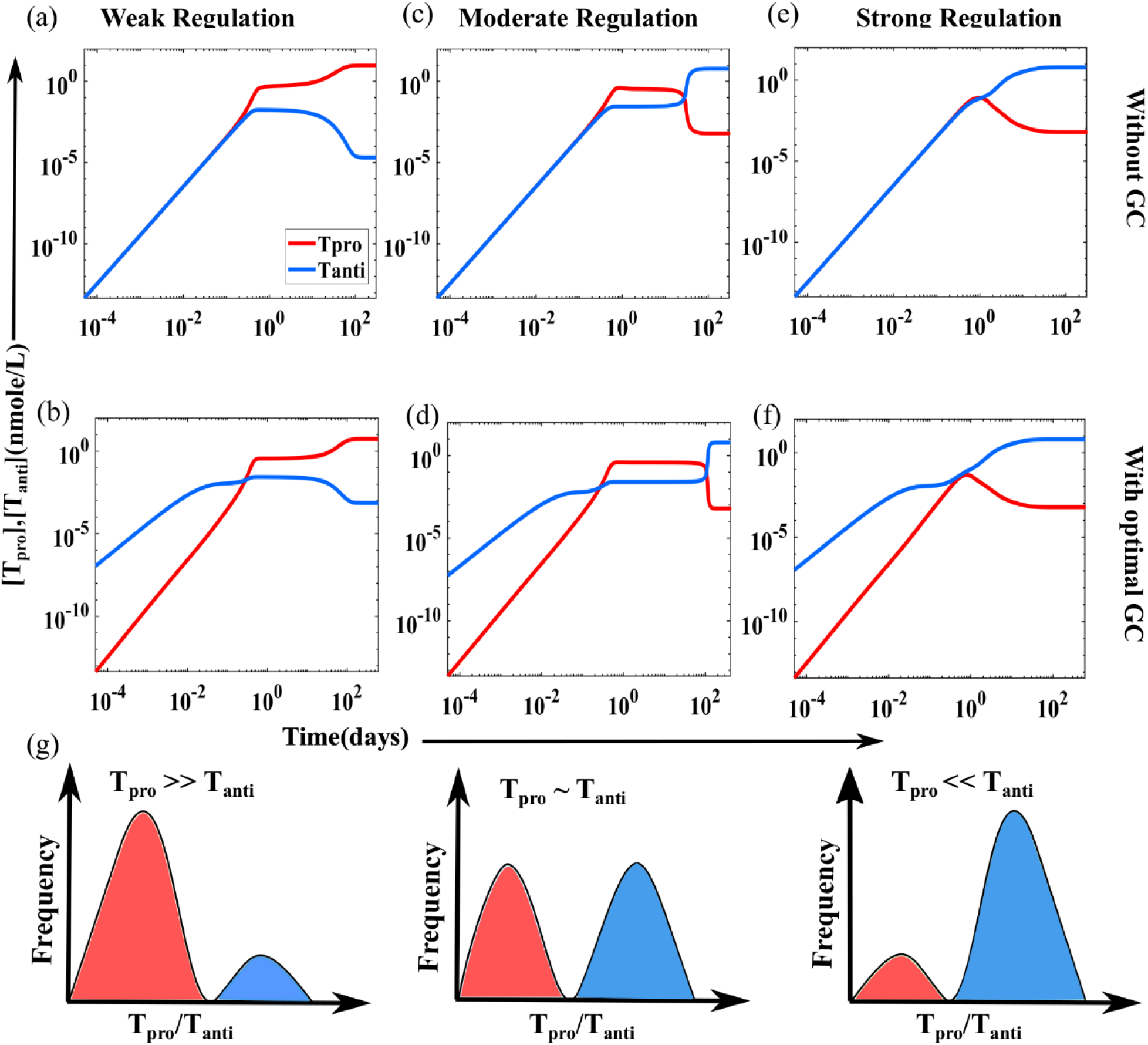
Time evolution of immune response showing all three regulations, both in the presence and absence of GC. A weak regulation state appears (a) in the absence of GC at *k*_*pro*_ = 50 and (b) in the presence of GC at *k*_*pro*_ = 70. Moderate regulation state appears (c) in the absence of GC at *k*_*pro*_ = 30, where the latent period is 20 days. (d) In the presence of GC Moderate regulation appears at *k*_*pro*_ = 56, where the latent period is 90 days. The system falls into a strongly regulated state at both (e) in the absence of GC and (f) in the presence of GC. The strong regulation remains strong both in the presence and absence of GC at the same value of *k*_*pro*_ = 10. As we vary *k*_*pro*_, other rate parameter values are kept constant and are taken from Table-I. A standard dose of Dex(GC) is taken to be optimal, which is 38.21 nmol/L (∼0.75 mg) [61]. For detailed explaination for determination of this optimal value (see Appendix B). In the plot, pro-inflammatory T cells and anti-inflammatory T cells are represented by the red line and the blue line, respectively. (g) We have shown a frequency distribution histogram of pro-inflammatory T cells(T_pro_) and anti-inflammatory T cells (T_anti_).

However, we find a shift in *k*_*pro*_, parameter value for strong and moderate regulations, when we change our system from the absence of GC to the presence of GC. We find that in the absence of GC, the system is under a strong regulation limit with *k*_*pro*_ = 50. However, when we introduce GC to our system at *k*_*pro*_ = 50, we find a moderate regulation which is shown in Fig. S4. The moderate regulation is extended over a wide range of limits, which is shown in Fig. S5. Beyond a certain limit, it falls in a weakly regulated regime. As presented in Fig. 3, The parameter values of *k*_*pro*_ are very sensitive for determining the strong regulation, moderate regulation, and weak regulation of T cells. Moreover, Fig. 3(g) shows a population distribution of number ratio between pro- and anti-inflammatory T-cells characterizing the overall classification weak, moderate, and strong regulation. This analysis was performed using Model-I.

### C. Intermediate stationary phase detection at moderate T-cell regulation limit

Using Model-II, we have replaced GC’s rate equation with a saturation function, which presents a one-time external dose intake mode. However, with an increase in the concentration of GC dose, the saturation function saturates into a constant value. Thus the coupling effect of GC is lost, shown in Fig. S7 of supplementary material. In our Model-I, we have accounted the natural metabolism of GC in the 5^th^ rate equation containing all the pharmacokinetic constants. This considers GC’s natural recursive regulation, which enables us to get a better understanding of the role of Glucocorticoid on T cell dynamics. Using a rate equation of GC makes our model more robust. We have observed a difference in the dose to obtain a moderate regime in Model-I and Model-II. For capturing the moderate regulation phase using saturation function, we need to go to the lower values of GC dose as exp(-G*) tends to zero with the increase in value of G*(dose of dex), which left us with loss of coupling effect of GC, so while using saturation function, we are constrained with lower GC dose. Despite the limitations of this saturation function, we used it for comparison purpose and to make the analysis more comprehensive with an existing model as used by Yakimchuk in a very recent work [33]. However, in both the methods, we have distinguished all three regulations: weak, moderate, and strong regulation. Most importantly, *the longer time dynamical analysis of T-cell population from any of these model studies at their moderate regulation limit consistently demonstrates the existence of an intermediate stationary phase behavior with a significant length of latent period as shown in Fig. 4*.

**FIG. 4:**
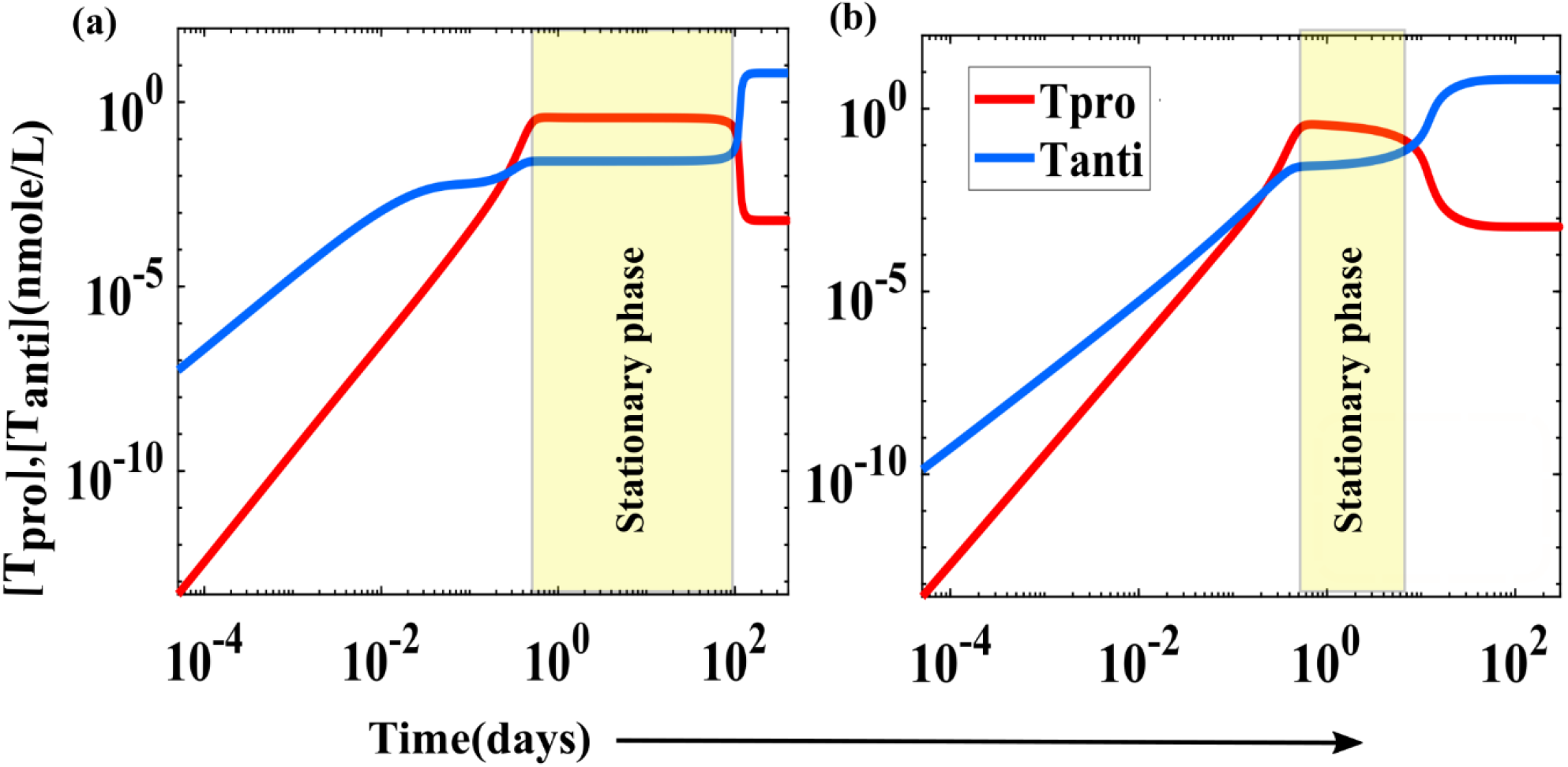
Time evolution of pro- and anti-inflammatory T cells in the presence of GC. (a) T-cell dynamics from Model-I, in the presence of standard/optimal concentration of GC=38.21 nM, (b) one-time external administration using saturation function GC=0.1 nM (∼1.96 microgram). In the plot, pro-inflammatory T cells and anti-inflammatory T cells are represented by the red line and the blue line, respectively. The yellow shaded region represents the latent period (stationary transition phase). Note that here we consider *k*_*pro*_=56, and the other rate values are the same as given in Table-I.

### D. Glucocorticoid and *k*_*pro*_ induced stationary phase optimization and immune phase diagram

In this work, GC is observed to have a significant role in efficiently keeping the immune system in a moderate regulation regime for a very longer duration of time. Moreover, it has also been seen that the moderate regime is conserved for a specific range of GC, which we consider an optimal range as shown in Fig. 5(a). Our findings also show that at a lower value or in the absence of GC, the latent period is small, leading to a risk of auto-immune disease due to the uncontrolled rapid growth of pro-inflammatory T cell (weak regulation). Moreover, when the GC dose is very high, the system again falls in the range of smaller latent period with significantly less population of pro-inflammatory T cells corresponding to an immune-compromised condition (strong regulation). In between strong and weak regulations, we observe the divergence of the system to a latent period peak, which corresponds to the moderate regulation. Our results signify the sensitivity of the immune system to the dose of GC. The optimal range of GC (Dex) is in accordance with the experimental finding [61,69] (for detailed explanation see Appendix B).

**FIG. 5:**
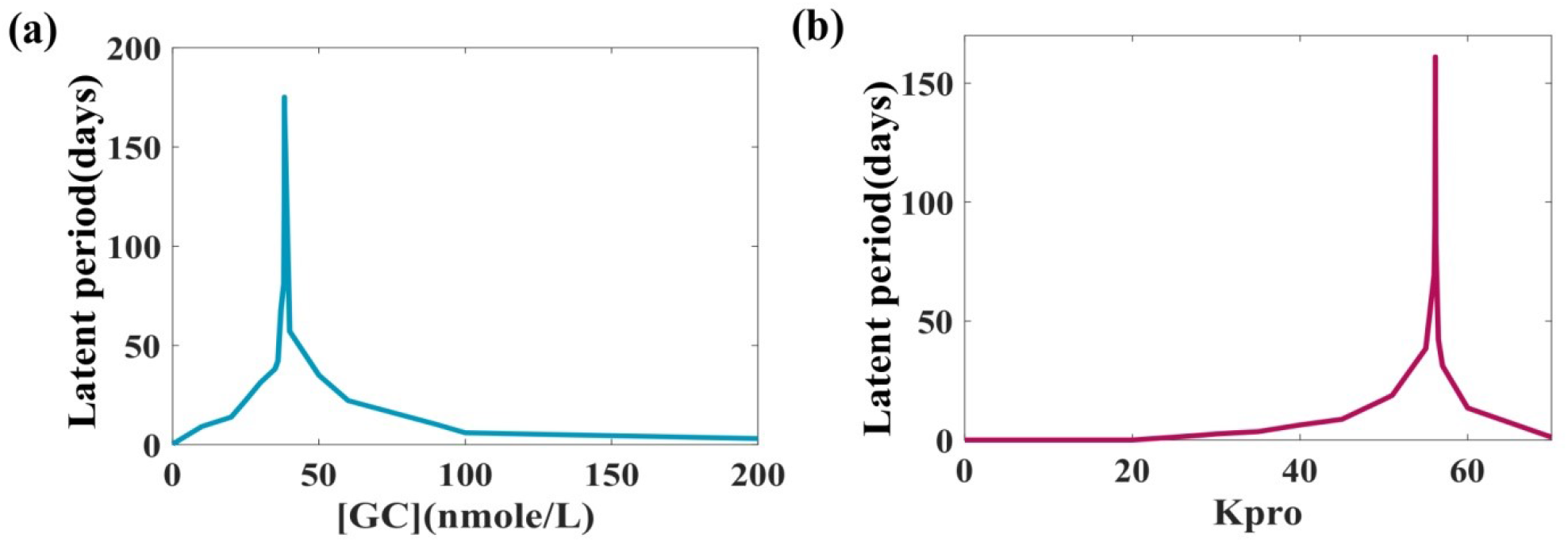
The sensitivity of the latent period to GC concentration and k_pro_. It is evident from the graphs that the system is very responsive to the dose of GC and *k*_*pro*_. Here we see a peak(divergence) in the latent period, which indicates that GC and *k*_*pro*_ induces moderate regulation of the immune system for a longer duration of time. (a)*k*_*pro*_=56.146 and other parameters/rate constants are taken from Table-I. The *k*_*pro*_ value is taken under moderate regulation. (b) As *k*_*pro*_ can vary from person to person, we have taken varied values of *k*_*pro*_ and determine the latent period, other parameters/rate constants are taken from Table-I, the dose of GC (Dex) is taken to be optimal which is 38.21 nmol/L the determination of this optimal value is explained in detail in Appendix B.

We have also looked into the sensitivity of the latent period with respect to the change of a sensitive rate constant, *k*_*pro*_, which is represented in Fig. 5(b). At a specific range of *k*_*pro*_ value, the immune system is in a moderate regulation with a large latent period. We have observed at both lower and higher values of *k*_*pro*._ The latent period is small, showing strong regulation and weak regulation, respectively. However, in a particular range of *k*_*pro*_ system between strong and weak regulation system fall is a moderately regulated regime. By changing *k*_*pro*_ values, we have modulated the system from strong regulation to moderate regulation to weak regulation. At *k*_*pro*_= 56.146, the system is at moderate regulation and has the largest range of latent period, and at *k*_*pro*_ =56.147, the system converts from moderate regulation to weak regulation, with a significant decrease in the latent period. This accounts for the sensitivity of *k*_*pro*_ value for its response in the latent period of immune system regulation.

As discussed above, the latent period is very sensitive response parameter to both GC and *k*_*pro*_, and with a change in the value of *k*_*pro*_ or GC we can observe a shift of the system’s regimes. However, our system in the presence of optimal doses of glucocorticoid shows the conservation of moderate regulation for large limits of *k*_*pro*_ and GC dose shown in Fig. S5 and Fig. S6, respectively in the supplementary material. As shown in Fig. 5, the latent period keeps increasing slowly, then there is sudden divergence, and it falls, modulating the system across weak to moderate to strong regulation limits. The modulation of latent period when it decreases can be an early warning signal of abrupt change in dynamics of the system from one stable state (moderate regulation) to another (weak regulation) upon small perturbation like small change in the GC dose or a sensitive rate parameter like, *k*_*pro*_ values.

With the classification of three regulation regions (weak, moderate, strong) we have investigated the boundaries between any two phases in the immune phase-space accounting for the above-mentioned two sensitive order parameters: *k*_*pro*_ and [GC]. The immune phase diagram is shown in Fig. 6. In light of our previous studies [3,4], it is worth mentioning here that distinguishing bistability in the moderate regulation regime is a key concept for understanding the basic phenomena of pro- and anti-inflammatory T-cell regulation. All these phenomena arise due to the nonlinearity of the biological system [70–72]. The phase diagram presented in Fig. 6 depicts the normalized concentration of pro-inflammatory T cell with respect to total CD4+ T cell concentration (sum of pro- and anti-inflammatory T cell) as a function of both *k*_*pro*_ and [GC]. It is plotted using numerical results from the solution of our system of equations obtained from Model-1. In this immune phase-diagram, the bistable/moderate regulation regime is a narrow constricted region. However, we find only near bistable regions; the immune system is sensitive to GC. At too weak/strong regulation, the system loses its GC sensitivity. It suggests that when a person is close to a moderate regime of parameter space, then only a synthetic GC intake may help.

**FIG. 6:**
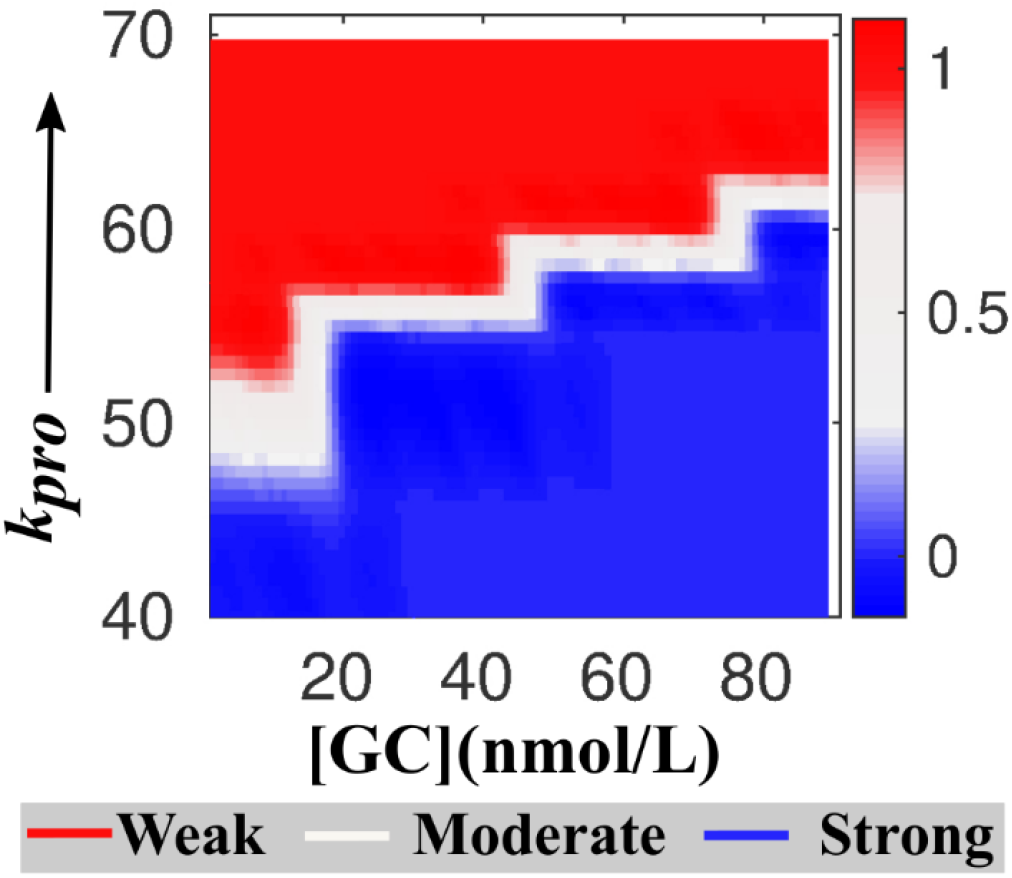
Steroid hormone, glucocorticoid induced immune phase diagram of CD4+ T cell. The graph depicts that the immune system is more sensitive to *k*_*pro*_ as a small change in the value of *k*_*pro*_ has potential to shift an immune regulation. However, GC sensitivity comes into action in the bistable/moderate regime. In the graph, the weak regime is denoted by redness, the strong regime is denoted as blueness, and the bistable/moderate regime is denoted as whiteness.

## IV. Conclusion

While the steroid hormone glucocorticoid (GC) is extensively used to control many acute and chronic inflammatory disorders, it is well documented that GC can cause a wide array of adverse effects those are both dose and time-dependent. Both the higher doses and long-term usage of low to moderate GC doses increase the risk of life-threatening infections due to the immunological cytokine imbalance and many associated factors [16,35,63,73,74]. While it is absolutely necessary to understand the immune responses of the T-lymphocytes, systematically, in a dose and time-dependent manner, clinically, it is a daunting task. However, a simple chemical dynamic model comprised of GC mediated cellular-level interactions amongst different subsets of T-cells and preliminary clinical guidance over known interaction-level database and parameters can provide valuable information about the state of our body. These approaches are often amenable to clinical measurements and certain conclusions. Thus, understanding the immune response of the complex human system requires collaboration between physicists, chemists, biologists, and clinical scientists. In particular, one may need to borrow the mathematical models and concepts from physics and combine certain rules of chemistry to understand the complex biological processes/interactions. There have been notable efforts in this interdisciplinary direction, although a lot remain to be achieved.

Below we summarized the key highlights of this study:

i. To monitor both time and dose-dependent GC effects on T-cell dynamics, we have developed two independent mathematical models. The first model considers the pharmacokinetic characteristics of GC, and in the other model; a saturation function is used to capture GC’s dose dependence of T-cell regulation. Both the models unanimously provide a similar dynamical pattern of pro and anti-inflammatory T-cells.
ii. In this long-term CD4+ T-cell kinetic study, three characteristic dynamic phases have been distinguished: growth (lag and log), stationary/latent period, and long-term-steady/memory phase. These phases are analogous to the growth kinetics of microbial cells.
iii. In this study, depending on the population ratio between the pro- and anti-inflammatory T-cell, we have quantitatively distinguished three classes of immune regulations: strong, weak, and moderate both in the presence and absence of steroid hormone, GC.
iv. A characteristic intermediate “stationary-phase” is detected to develop especially in the moderate regulation limit under the influence of pathogen. This is an apparent near-normal clinically asymptomatic/latent phase, and the corresponding latent period can sustain over for a long time (more than a couple of months), where pathogenic load drops dramatically. The emergence of prolonged clinical latency correlates well with the CD4+ T-cell dynamics that have been monitored in the case of HIV infection [32].
v. The study finds the latent period in the intermediate stationary phase to vary non-monotonically as a function of the concentration of steroid hormone, GC. At a standard/optimal level of GC concentration, the latent period is found to reach the peak implying that GC optimizes the stationary phase by subtly balancing the population ratio between the pro- and anti-inflammatory T-cell. However, this needs to be clinically verified.
vi. At a longer time, after the asymptomatic phase, a long-term steady-state outcome is reached, which is a decision making phase of the system. In this decision making phase, the system either deactivates or reactive the pro-inflammatory actions, which decides the fate of the antigen/disease. In the presence of steroid administration, after a prolonged stationary phase, the system has a general tendency to switch on a strong regulation limit where immunity is challenged. And, here comes the relevance of dose and time dependence of steroid drug administration. It is now well-known that in addition to GC, many other immunosuppressive drugs such as prednisone, pain-killers (morphine, codeine, hydrocodone, opioids), and other chemotherapeutic drugs perform life-saving tasks within human body but often causes pathogen/viral reactivation when immunity is critically suppressed. In fact, during chemotherapy, viral reactivation event is very real [75–77].
vii. Finally, understanding the final steady-state outcome in different immune regulation limits and their co-existence are illustrated by an immune phase-diagram which reveals that the steroid-dependent moderate/healthy phase is a constricted immune regime which is adept at tolerating a wide range of pathogenic stimuli when the auto-catalytic rate of pro-inflammation is low (i.e., when T-cells are less aggressive).

While the immunosuppressive drug, GC potentially controls weak/autoimmune-prone T-cell regulation, the study reveals a new odd characteristic of steroid-dependent T-cell dynamics, which highlights a prolonged stationary phase. In the stationary phase, while the system is auto-immune controlled under the steroid treatment, it is now more vulnerable to pathogen reactivation. These challenges can be circumvented by long-term diagnostic measures, especially to monitor the prolonged latency in the intermediate stationary phase and by prescribing antiviral treatment along with GC treatment in an effort to ward off pathogen reactivation as suggested by few clinical studies [32,78,79]. Our study indeed provides the rationale behind such mode of treatments and encourages the awareness against imprudent steroid medication.

In this study, we have employed a simple chemical network model of the immune system to understand the effect of steroid drug like GC. This eventually results in an optimized stationary phase to control auto-immune disorders without any adverse effect at least for some time (if not longer). The non-monotonic steroid dependence of the intermediate stationary phase can be used as a diagnostic marker for steroid treatment, especially when the system loses bistability. In future, we shall attempt to explore more towards the noise-induced bistability and fluctuation driven immune response to monitor how the fluctuation of certain T-cell subset affects the stationary phase latency and steady-state outcome during steroid treatment.

## Supplementary Material

See supplementary material to follow the description of course of evolution of glucocorticoid as a drug, along with a large complex network used for coarse-grained network model development, parameter estimation, detailed steady-state and stability analysis, and other data analysis.

## Acknowledgments

This work was supported in parts by start-up grants from IISER Kolkata. SR acknowledges support from DBT, India (Grant No. BT/12/IYBA/2019/12).

## Appendix A Simulations in the absence of Glucocorticoid

In order to look at the dynamics of the system in the absence of Glucocorticoids we remove all the terms involving the glucocorticoids. Hence, we would have four coupled ODE, as given below.

### Coupled ODE for the system in the absence of Glucocorticoid (representing drug-free immune regulation)

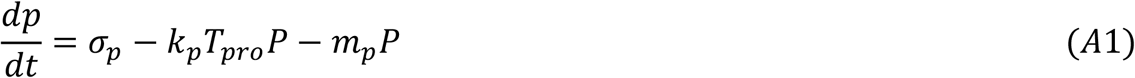

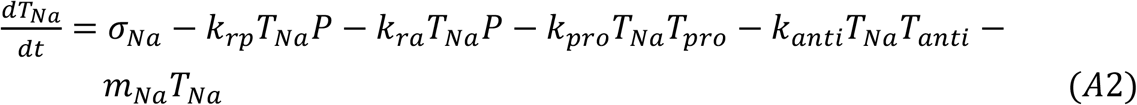

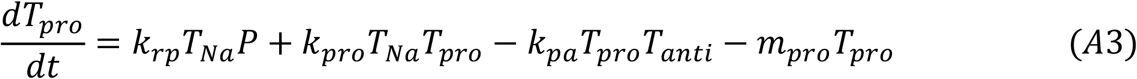

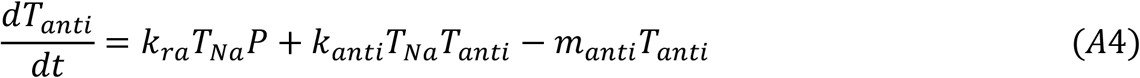

## Appendix B Parameter estimation and other calculations

### Determination of initial Naïve T cell concentration

We have assumed that in the absence of antigen, hundred CD4+ naive T-cells can pre-exist within this fixed volume (100 nano-liter) [84].

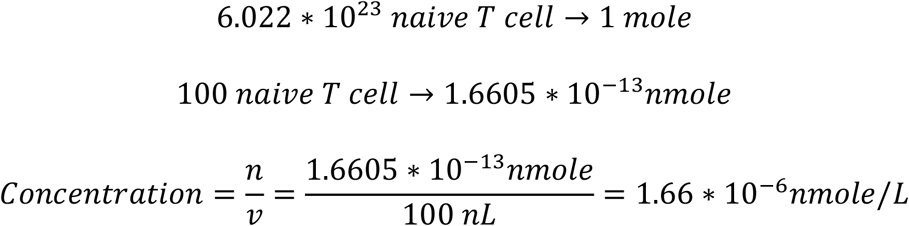

### Determination of GC dose

The dose of Dexamethasone at which it shows its immunomodulatory activity is 0.75 mg [61,69].

The volume Distribution is Dexamethasone is 40-60 L [61,69].

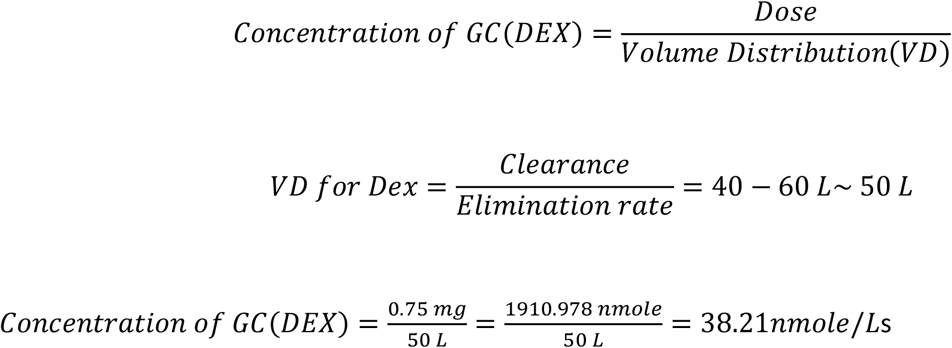

### Parameter Estimation

In the work of Fouchet et al., they have considered APC activation by antigens, effector T cells, and regulatory T cells to be variable [61,69]. However, they have considered the naïve T cell maturation rate activated by APC to be 1. So, in our work, we have considered the following values range from 0.1 to 0.01. Based from our early study [3,4], as the antigen of the invaded pathogen or self-antigen leads to activation of Resting APC and Active APC, which further leads to activation of the maturation process of Naïve T cells into Pro-inflammatory T cells and Anti-inflammatory T cells from which, we can conclude that the antigen-induced rate of differentiation of naïve T cell to the pro-inflammatory and anti-inflammatory T cell, can be the sum up of APC activation by antigens and naïve T cell maturation rate activated by APC.

However, the Glucocorticoid related pharmacokinetic rate constants are taken from various literatures [60,61,69,80,81].

Absorption rate of Glucocorticoid(dex) [61] = 4.8729±8.4998 1/hour Bioavailability of Glucocorticoid(dex) [60] = 70-78%(75%) Biological half life of Glucocorticoid(dex) [61] = 36-72 hour(40 hour) Rate of decay of Glucocorticoid(dex) = *m*_*G*_*

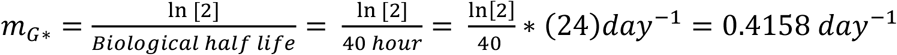

## Appendix C Steady-state and stability Analysis

### For the system to be in a steady-state, it should be the following these conditions

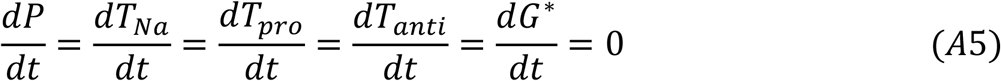

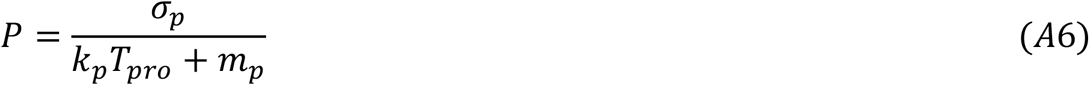

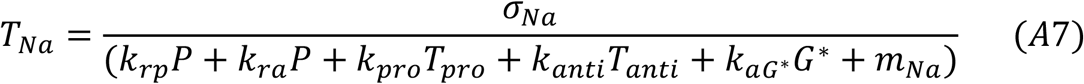

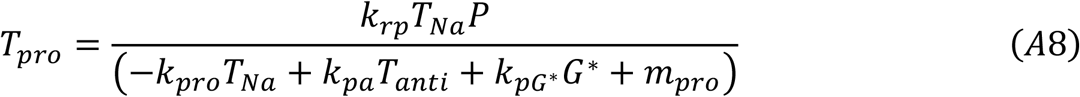

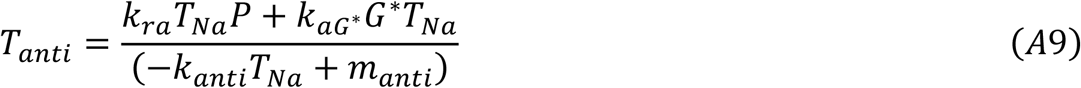

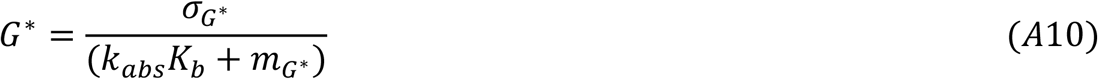

The latent period and steady-state found from our MATLAB plot satisfy these steady-state conditions.

### The Jacobian matrix for the given system of ODEs (1-5) represented as

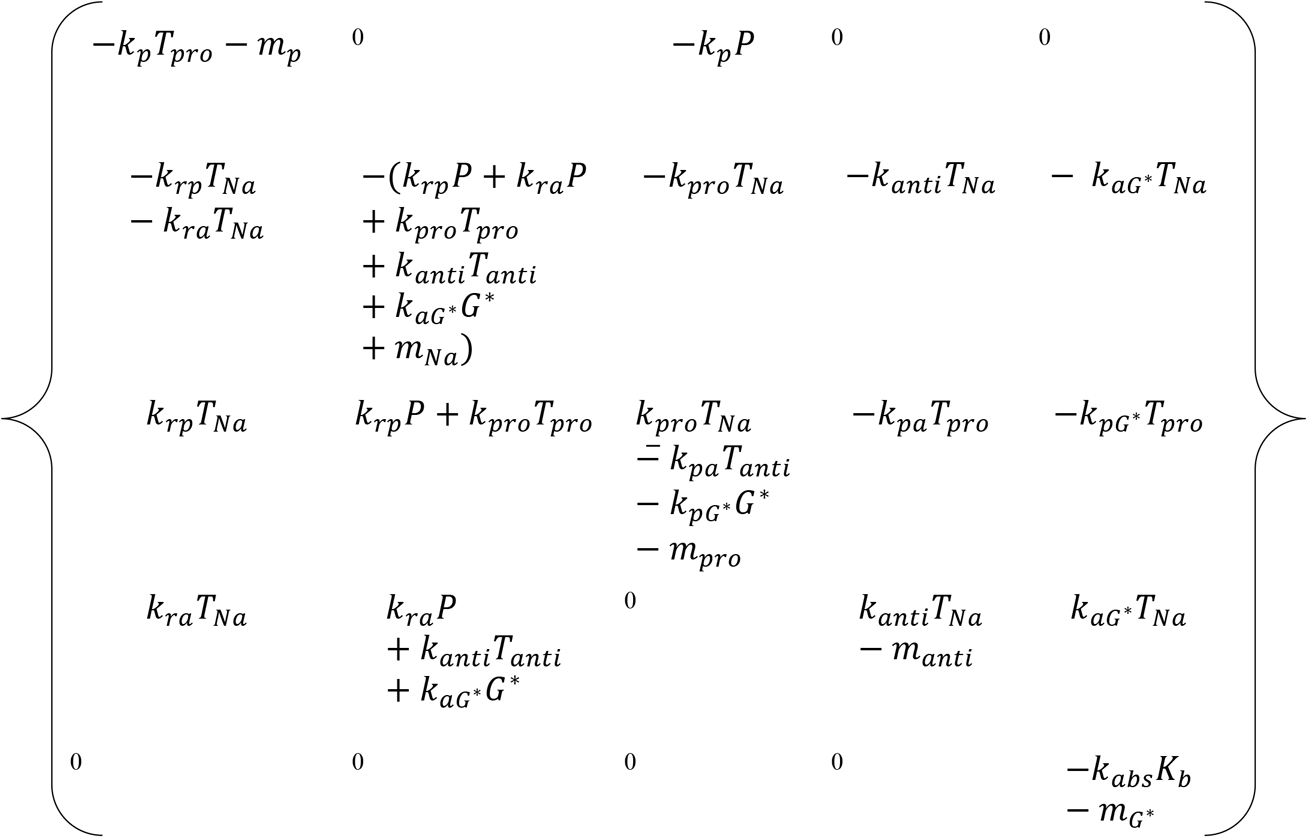

To analyze the steady-state conditions when there are no source parameters. There is an equilibrium point when all the concentration of all the element is equal to zero:

*E*(*P, T*_*Na*_, *T*_*N*_, *T*_*aN*_, *G*^*^) = (0,0,0,0,0). Then Jacobean matrix is presented as:

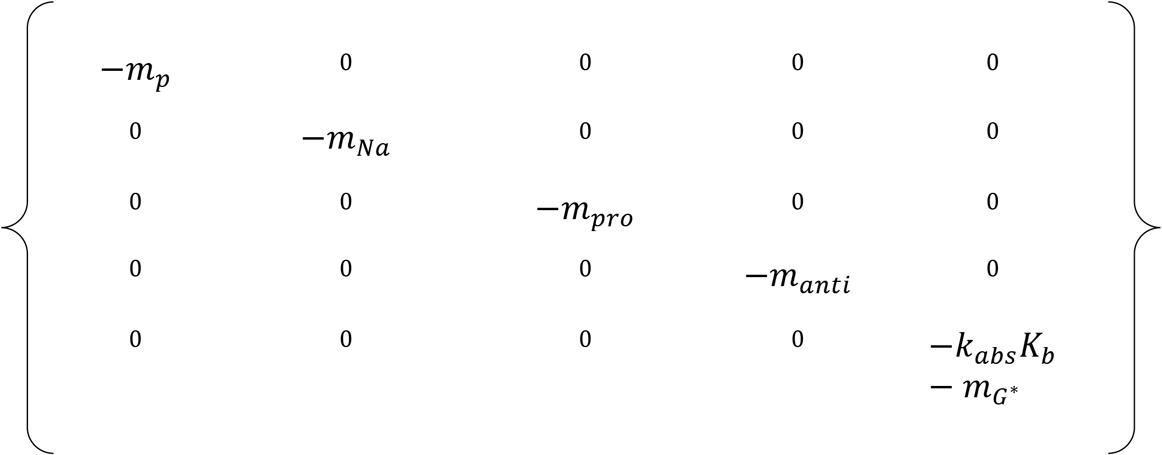

Then the eigenvalues of the Jacobean matrix are

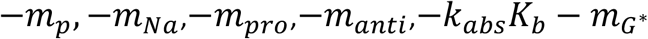

Since all the rate constants considered in the system are positive. This concludes the five eigenvalues of the system is negative, which justifies the stability of the considered system.

### Saturation point for respective ODE when we consider each ode is mutually exclusive

From Eq (A6) P will saturate at

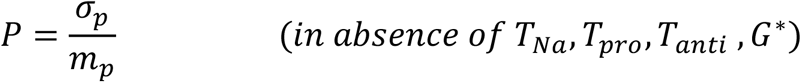

From Eq (A7) *T*_*Na*_ will saturate at

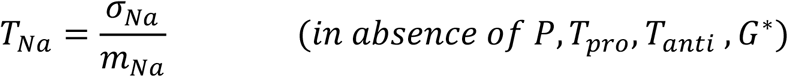

From Eq (A8) *T*_*N*_will saturate at

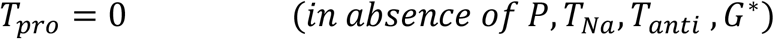

From Eq (A9) *T*_*aN*_will saturate at

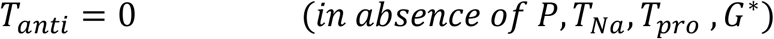

From Eq (A10) *G*^*^ will saturate at

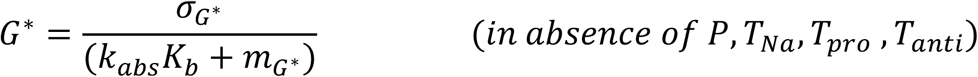

## Data Availability Statement

The data that support the findings of this study are available in the main text and supplementary material but can also available from the corresponding author upon request.

## Notes

### Competing Interest Statement

The authors have declared no competing interest.

