## Supplementary Information for "Immune phase transition under steroid treatment"

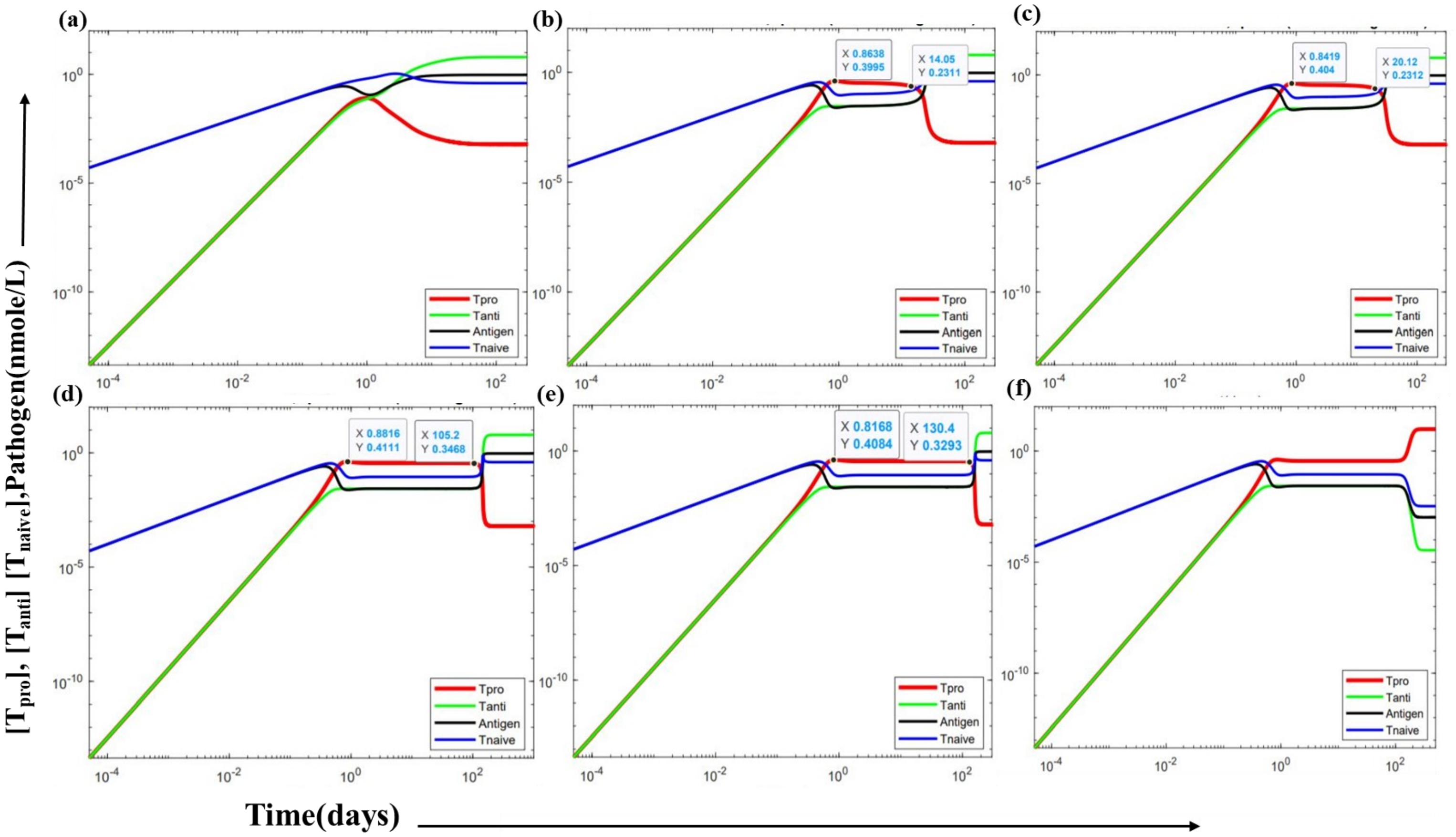


**Figure S1: Time evolution of immune response in absence of GC.** Time evolution of Pro-inflammatory T cells, Anti-inflammatory T cells, Naïve T cells and Pathogen are plotted by solving coupled kinetic equations shown in **Text S3** using a deterministic approach in MATLAB. (a)$k_{pro}$=10, (b)$k_{pro}$=29 (c)$k_{pro}$=30, (d)$k_{pro}$=31.2327, (e)$k_{pro}$=31.23279 (f)$k_{pro}$=31.2328. As depicted by the graphs $k_{pro}$ parameter values are very sensitive and for determining the strong regulation, moderate regulation, and weak regulation of pro-inflammatory T cell(other parameter values are taken from **Table 1**). Note plot obtained using Model I.

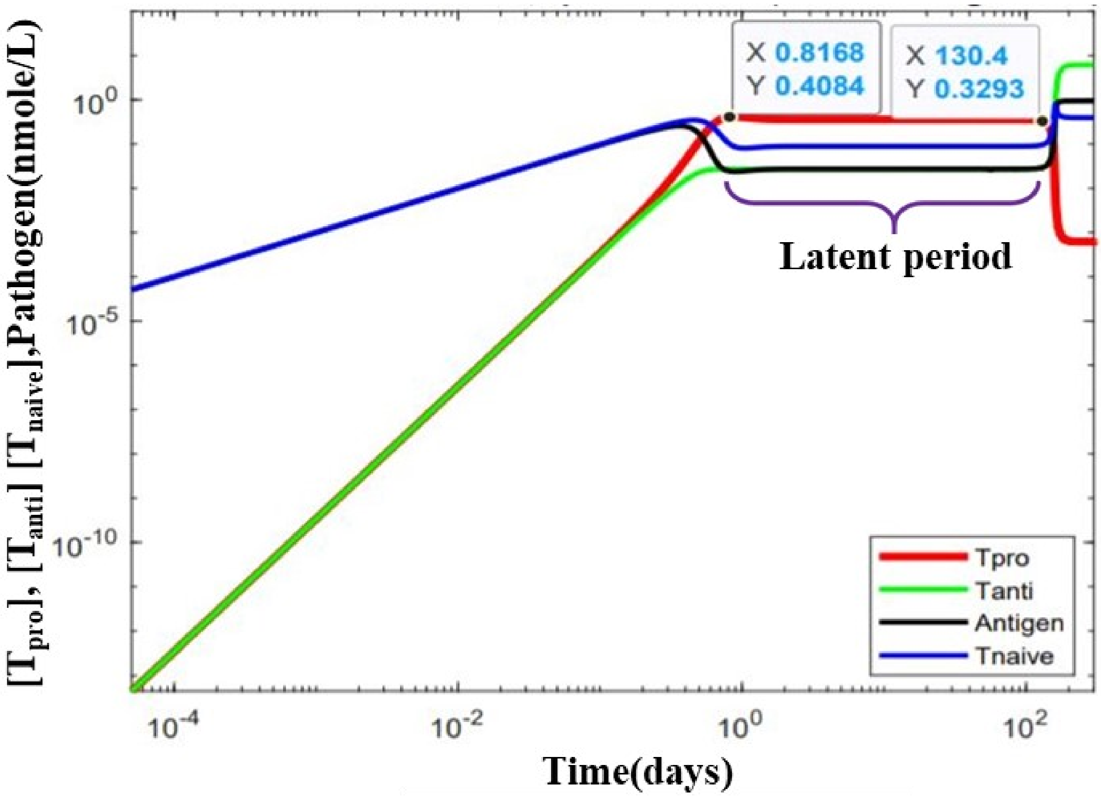


**Figure S2: Representation of the latent period.** The time range within which the concentration of all the cell population (Pro-inflammatory T cells, Anti-inflammatory T cells, Naïve T cells, and pathogen) are not changing with time, and after that, there is a jump to the steady-state condition. This graph is plotted in the absence of GC at $k_{pro}$ =31.23279, and other rate constants are the same as given in Table 1. It is evident from the plot that the latent period for these particular rate constants is around 129.5832 days(130 days approx). Note plot obtained using Model I.

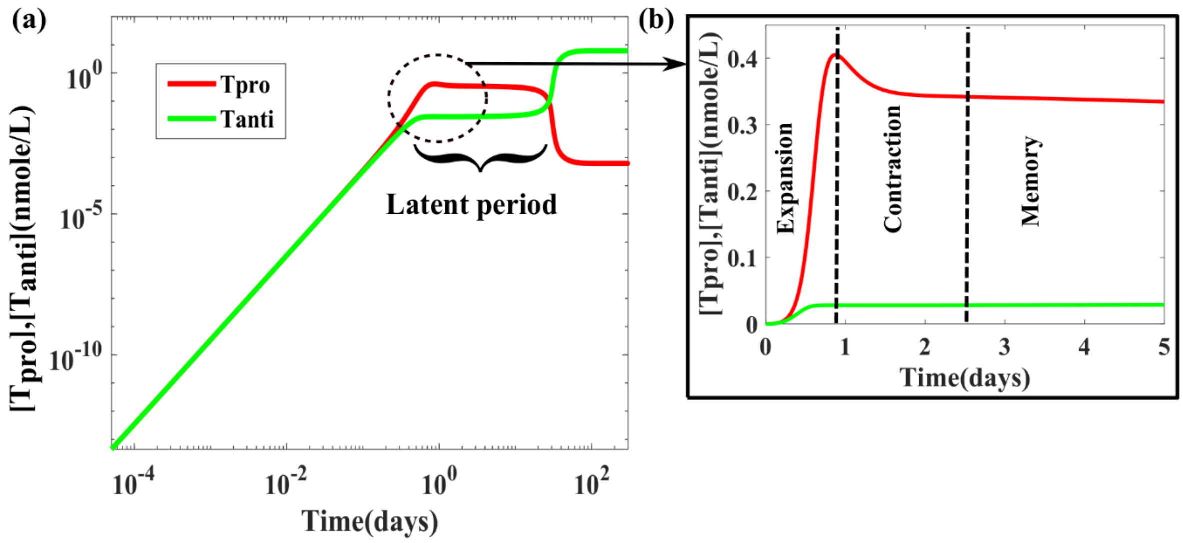


**Figure S3: Time evolution of the immune response of CD4+ T cells in the absence of any drug.** The plot represents a regime of the immune phase regulation that contains a latent period, i.e., the time range within which the concentration of Pro-inflammatory T cells and Anti-inflammatory T cells does not change with time. After the latent period, there is a jump to the steady-state condition. Moreover, the schematic illustration of the phases of immune response mediated by antigen-specific Pro-inflammatory T cells is depicted on the left side where three phases of the Pro-inflammatory T cells immune response (expansion, contraction, and memory) are indicated. The time evolution of Pro-inflammatory T cells and Anti-inflammatory T cells are plotted by solving coupled kinetic equations in MATLAB using a deterministic approach. In the plot, Pro-inflammatory T cells and Anti-inflammatory T cells are represented by the red line and the green line, respectively. Note that here we consider $k_{pro}$ =30, and the other rate values are the same as given in Table 1. Note the zoomed portion of (a) is shown in (b). Note plot obtained using Model I.

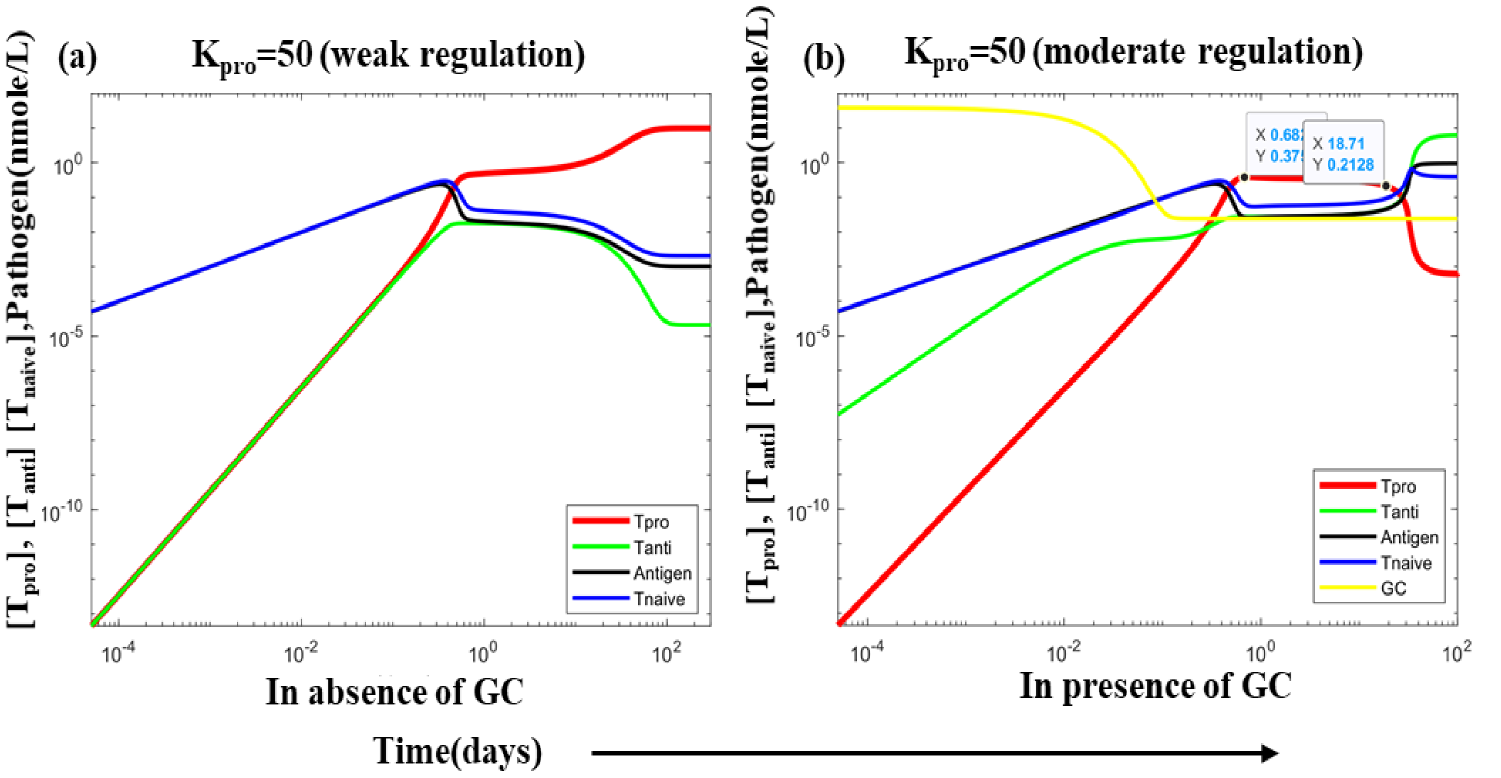


**Figure S4: Comparing the shift of regime when Glucocorticoid is introduced in the system**, the weak regulation shift to moderate regulation at the same $k_{pro}$ value of 50. A weak regulation state appears at (c) in the absence of GC at $k_{pro}$ =50. (f) In the presence of GC at $k_{pro}$ =50, we found a moderate regulation. Note plot obtained using Model I.

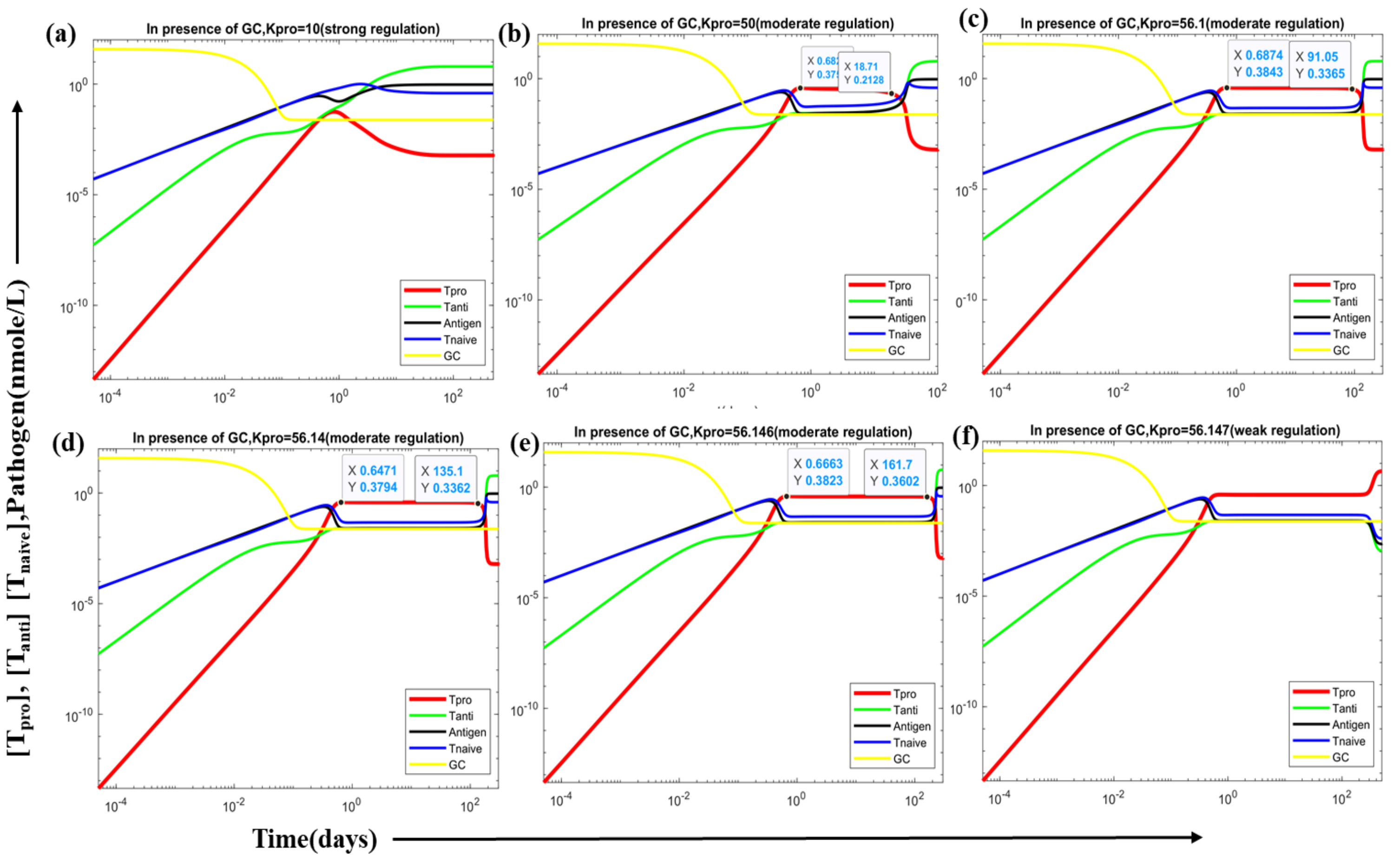


**Figure S5: Representation of modulation of regulation and response function(latent period) with different values of $k_{pro}$ in the presence of an optimal dose of GC.** With variation in $k_{pro}$, we have found that the latent period increases and becomes largest at (e) $k_{pro} = 56.146$, representing a system with moderate regulation and (f) $k_{pro} = 56.147$, the system falls in weak regulation, after falling in weak regulation latent period keep on decreasing. Before the regime shift from moderate to weak, we have observed a critical slowing down of the system(latent period becomes approximately 160 days at $k_{pro} = 56.146$), indicating an early warning signal before regime shift. (a) latent period= 0 days, $k_{pro} = 10$ (b) latent period= 18.03 days, $k_{pro} = 50$, (c) latent period= 90.3626 days, $k_{pro} = 56.1$, (d)latent period= 134.45 days, $k_{pro} = 56.14$, (e) latent period= 161.033 days, $k_{pro} = 56.146$. Apart from $k_{pro}$, other parameter values are taken from **Table 1**. The initial dose of Dex(GC) is taken to be optimal, which is 38.21 nmol/L. Note plot obtained using Model I.

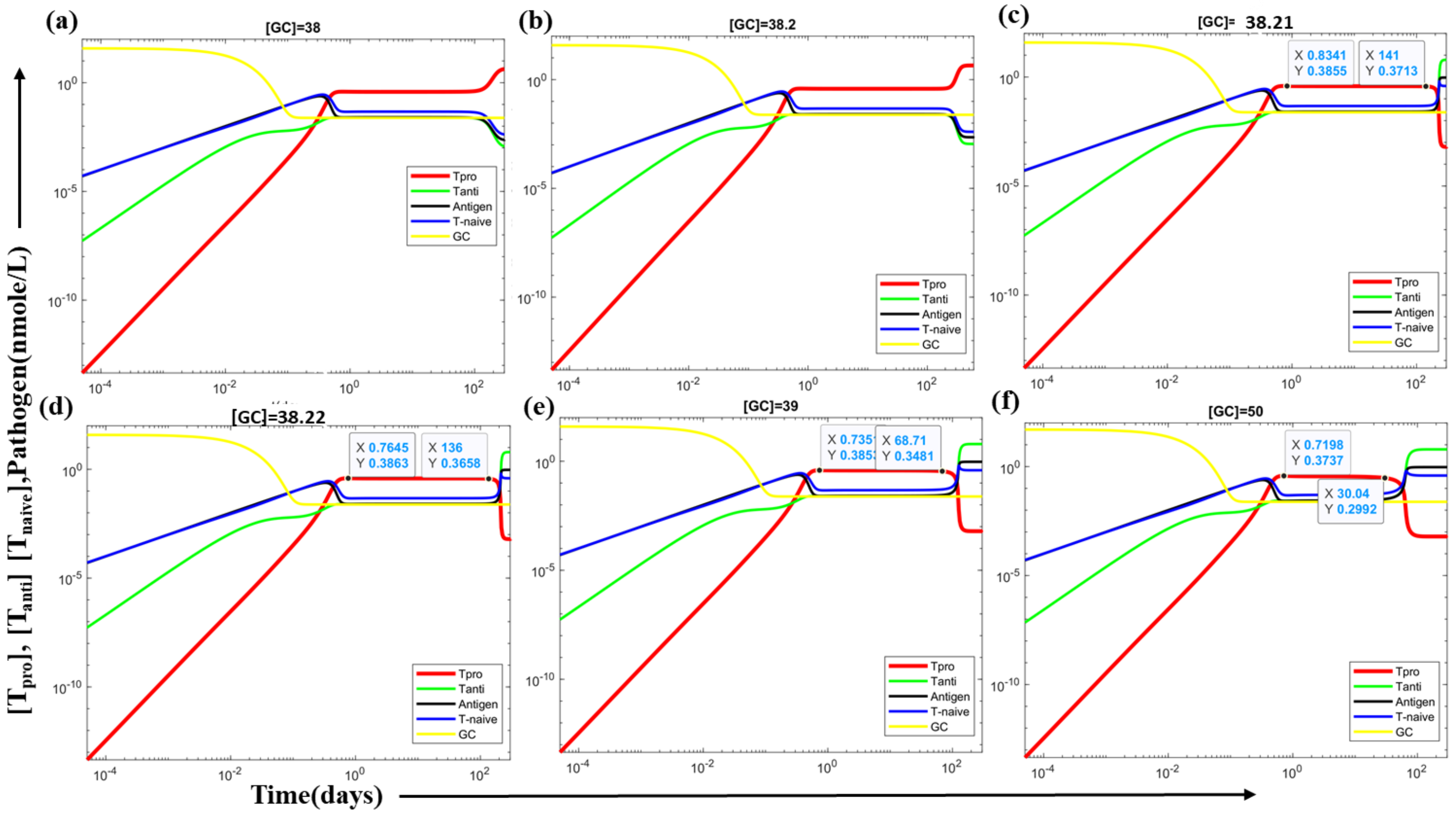


**Figure S6: Representation of modulation of regulation and response function(latent period) with different values of the initial dose of GC.** We have found upon varying the initial pre-existing dose of GC, the latent period increases and become largest at (c) GC= 38.21 nM representing a system with moderate regulation and (b) GC= 38.2 nM, the system falls in weak regulation, after falling in weak regulation latent period keep on decreasing. Before the regime shift from moderate to weak, we have observed a critical slowing down of the system(latent period becomes approximately 140 days at GC=38.21 nM), indicating an early warning signal before regime shift. Here we have taken $k_{pro}$=56.416; other parameter values are taken from **Table 1**. Note plot obtained using Model I.

**The outcome of replacing rate equation of GC by a saturation function (method II)**

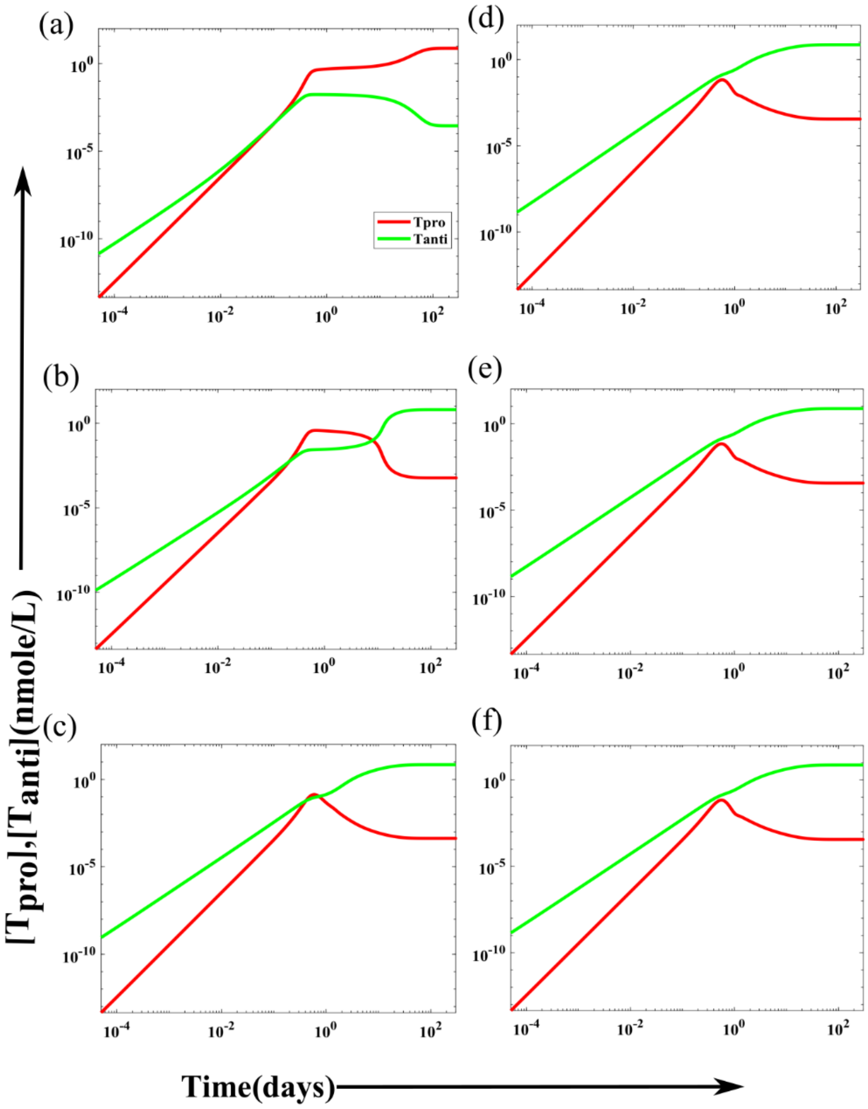


**Figure S7: Time evolution of immune response using saturation function showing all three regulations, in the presence of different value of GC.** A weak regulation state appears at (a) In the absence of GC= 0.01 nM. (b) In the presence of GC=0.1 nM Moderate regulation, state appears. However, a system falls to a strongly regulated state at (c)GC=1 nM, (d)GC=10 nM, (e)GC=50 nM, (f)GC=80 nM. Here we have taken $k_{pro}$=56, and other parameter values are taken from **Table 1**. Note that: (0.1 nM=1.96 microgram). Note plot obtained using Model II.
